# A Human Neuronal Co-Culture System Reveals Early Tumor-Neuron Communication and Targetable Pathways in Glioblastoma

**DOI:** 10.1101/2025.09.10.675396

**Authors:** Ouada Nebie, Niyi Adelakun, Liwen Zhang, Luke Kollin, Brian Fries, Akhil Medikonda, Monica Venere, Pierre Giglio, Nam Chu, Nhat Le

## Abstract

**Background:** Glioblastoma (GB, IDH wild type) is the most aggressive primary brain tumor in adults, with recurrence driven by residual tumor cells that re-establish interaction with surrounding neurons. While neuronal activity is recognized as a driver of GB progression, the earliest neuronal responses to tumor contact remain poorly understood. Existing models rarely capture these acute events or the heterogeneity of responses generated by tumors of different origins, limiting insight into the earliest neuron-tumor interactions.

**Methods:** We developed a dual-interface human iPSC-derived neuronal culture system to investigate acute neuronal responses to glioblastoma exposure. Neurons were challenged with either established GB cell lines or patient-derived glioblastoma cells (PDGCs). Using quantitative proteomics, high-resolution imaging, and immunological assays, we characterized compartment-specific neuronal changes and mapped activated signaling pathways. We also screened selective inhibitors for their effects on both tumor proliferation and neuronal integrity.

**Findings:** Within 24 hours of exposure, neurons displayed synaptic remodeling and activation of GB-related signaling cascades. Proteomic analysis of GB exposed neurons revealed enrichment of pathways associated with GB and abnormalities in neuronal circuits. Notably, the U-87MG cell line, but not PDGCs, induced pronounced synaptic disruption, neurite retraction, and MAPK pathway activation, with distinct molecular signatures across neuronal compartments. ERK1/2 and p38 MAPK signaling were differentially activated depending on the GB cells source, correlating with specific structural and functional synaptic alterations. Targeted inhibition of MAPK components significantly suppressed U-87MG proliferation and preserved neuronal architecture.

**Interpretation:** We present a human neuronal culture model that detects and discriminates the earliest neuron-derived responses to glioblastoma from diverse tumor sources. By linking neuronal remodeling to tumor-specific signaling pathways, the platform uncovers both the heterogeneity of neuron-tumor interactions and early, targetable vulnerabilities. This system offers a translational tool to advance understanding of GB recurrence and to guide development of therapies with dual neuroprotective and anti-tumor efficacy.

**In Brief:** Early interactions between glioblastoma and human neurons drive rapid, source-specific synaptic remodeling mediated by compartmental MAPK pathway activation. This neuronal co-culture model identifies distinct profiles of tumor-neuron communication, highlights synaptic vulnerability as a therapeutic axis, and demonstrates that MAPK pathway inhibition yields both neuroprotective and anti-tumor effects.

**Summary:** Glioblastoma (GB) co-opts neuronal circuits to drive tumor progression, yet the earliest neuronal responses that may shape recurrence remain poorly defined. We developed a human iPSC-derived neuronal co-culture model that captures acute communication between neurons and glioblastoma from diverse sources, including serum-adapted cell lines and patient-derived cells. Within 24 hours, glioma-exposed neurons exhibited synaptic remodeling and activation of tumor-associated signaling pathways. High-resolution imaging and proteomics revealed compartment-specific synaptic alterations, with ERK1/2 and p38 MAPK signaling differentially engaged depending on the tumor source, corresponding to distinct structural and functional outcomes. Pharmacologic inhibition of MAPK components both suppressed tumor cells growth and preserved neuronal integrity. By modeling source-dependent and early neuron-tumor interactions, this platform not only identifies MAPK signaling as a critical mediator of synaptic vulnerability but also provides a clinically relevant tool for investigating the mechanisms of glioblastoma recurrence. It offers a framework for developing therapies that target the dual vulnerabilities of tumor progression and circuit remodeling.

Highlights

- A dual-interface human iPSC-derived neuronal co-culture system models early neuron-glioblastoma (GB) interaction.
- Serum-adapted U-87MG, but not serum-free patient-derived GB cells, induces pronounced synaptic disruption and neurite retraction.
- Global proteome of neurons reveals GB-associated signatures and neuronal circuit alterations in response to both U-87MG and patient-derived GB cells.
- ERK1/2 and p38 MAPK signaling are differentially activated in neuronal compartments, depending on GB source.
- MAPK pathway inhibition suppresses U-87MG proliferation and preserves neuronal integrity, revealing actionable neuroprotective and anti-tumor targets.

## INTRODUCTION

Gliomas are primary brain tumors that arise from glial cells, with glioblastoma (GB) representing the most aggressive and lethal subtype. Despite advances in surgery, radiotherapy, and chemotherapy, GB remains uniformly fatal, with a median survival of just 15 months [1–3]. Recurrence, rather than initial tumor growth, is the ultimate cause of death, yet its biological drivers remain unknown [4]. This poor outcome reflects not only the tumor’s resistance to therapy but also its ability to remodel the brain microenvironment, including neighboring neurons, to facilitate growth and invasion [5–7]. Recent studies have revealed that neuronal activity actively contributes to glioma progression. GB cells form synaptic-like connections with neurons [8–11] and co-opt excitatory neurotransmission to drive tumor growth and treatment resistance [12, 13]. These interactions also disrupt normal neuronal function, enhancing excitotoxicity and altering circuit dynamics [14]. Paradoxically, some standard therapies, including radiation and anti-angiogenic agents, may further promote infiltration into healthy neural tissue [15, 16]. However, modulating neuronal activity has been shown to slow tumor growth [17–19], underscoring the importance of understanding glioma-neuron interactions early in the disease progression.

Despite this emerging interest, mechanistic insight into how glioma cells influence human neurons during initial tumor infiltration remains limited. Existing animal models and human in vitro systems lack the resolution or physiological relevance needed to examine acute, spatially defined tumor-neuron dynamics [20, 21]. Additionally, most therapeutic development still focuses primarily on tumor-intrinsic targets, often overlooking the tumor-neuron interface, which is increasingly recognized as a driver of recurrence, disease progression and therapeutic resistance [14, 22–25].

To address this, we created a co-culture platform derived from human iPSCs to investigate early interactions between glioma and neurons. Using both U-87MG and patient-derived glioblastoma (GB) cells, we discovered that GB exposure promptly triggers synaptic remodeling and MAPK pathway activation in neurons, driving them into a tumor-promoting state. MEK inhibition protected neurons while suppressing glioma proliferation/invasion. This model allows for high-resolution investigation of tumor-neuron interactions. This streamlined model system not only facilitates rapid, physiologically relevant analysis of tumor-neuron dynamics, but also serves as a powerful platform for dissecting the molecular mechanisms of glioma-induced synaptic vulnerability and testing candidate therapeutics at early stages of tumor-neuron interaction.

## RESULTS

### 1. A human dual-interface co-culture system to study early neuronal response to glioblastoma

To investigate the early molecular and functional effects of glioblastoma on human neurons, we established a dual-interface co-culture system using GB cells and human iPSC-derived cortical neurons (Fig. 1A). This system was specifically designed to model the acute phase of tumor-neuron interactions and overcome the limitations of static, non-compartmentalized culture systems. Unlike traditional organoids or mixed co-cultures that lack spatial precision, our approach allows direct yet controllable exposure of neurons to GB cells, enabling the identification of rapid, cell-type-specific changes at the tumor-neuron interface.

**Figure 1.**
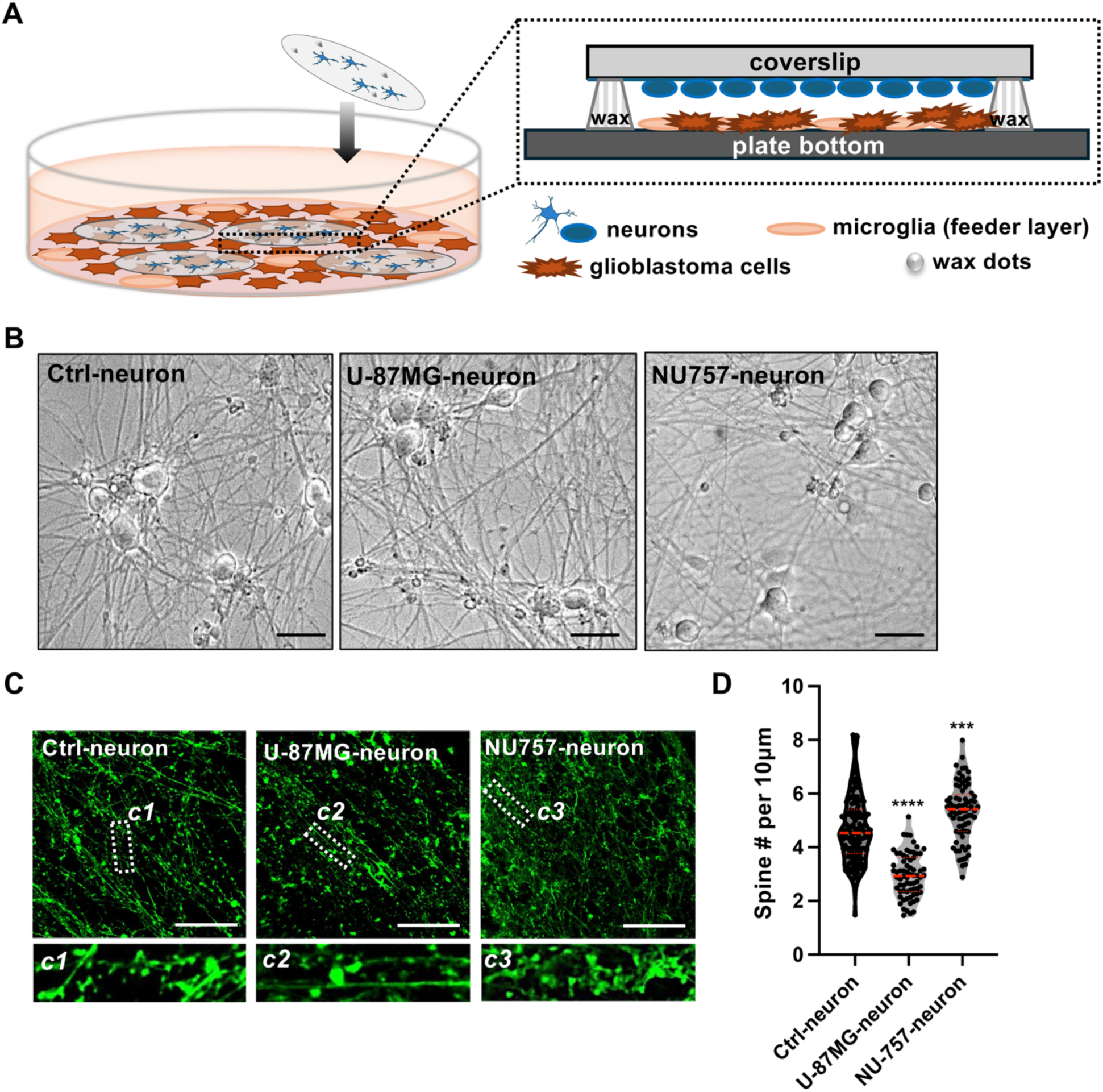
A dual-interface human glioblastoma–neuronal co-culture system reveals glioma-induced changes in neuronal morphology. **(A)** Schematic of the dual-interface human glioblastoma (GB)-neuronal co-culture system used to study acute neuronal responses to glioma. Human iPSC-derived neurons are cultured on glass coverslips suspended approximately 0.5 mm above a layer of human microglial cells using wax dots as spacers. GB cells are introduced into the microglial layer beneath the neurons. **(B)** Representative differential interference contrast (DIC) images of human iPSC-derived neurons cultured under three conditions: control (Ctrl-neuron), exposed to U-87 MG cells (U-87 MG-neuron), or exposed to serum-free patient-derived NU-757 cells (NU-757-neuron) at 24 hours. Scale bars = 30 µm **(C)** Fluorescent phalloidin staining (green) of dendritic spines in neurons after 35 days of differentiation under the same conditions as in (B). Insets (c1, c2, c3) show magnified views of the dashed regions. Scale bars = 50 µm. **(D)** Quantification of dendritic spine density (spines per 10 µm) corresponding to (C). Data were pooled from 70-160 dendritic segments across 2-3 independent experiments. U-87 MG exposure significantly reduced spine density, whereas NU-757 exposure led to increased spine density compared to controls. Data are shown as mean ± SD. Statistical comparisons were performed using unpaired t-tests. Significance: ***P < 0.001; ****P < 0.0001.

We employed a neuronal differentiation protocol [26] in which human iPSCs (hiPSCs) can be induced to differentiated cortical neurons in an efficient and scalable manner. Human iPSCs were differentiated into excitatory cortical neurons over 3-4 weeks and matured in monolayer culture on coated coverslips. Neuronal identity and synaptic maturity [27] were validated through expression of MAP2, synaptophysin, and PSD-95 (Fig. S1). Neurons differentiated from hiPSCs develop functionally active synapses [26]. On day 35 of post-differentiation, neurons were suspended above a human microglia layer and maintained in serum-free medium. GB cells were introduced into the system by being seeded onto the bottom microglia layer to allow molecular exchange without overgrowth (Fig. 1A). This configuration enabled acute co-culture periods, allowing glioma-derived factors to influence neurons before irreversible structural changes or tumor overtake occurred.

To ensure the system maintained biological relevance during the acute exposure window, we confirmed neuronal viability and morphologies in control condition without exposure to glioblastoma. Neurons in these cultures, which are maintained up to weeks in the presence of a human microglia cell layer, keep morphologically matured axons and dendrites (Fig. 1B, Ctrl-neuron), and functional excitatory and inhibitory synapses [28].

The dendrites are studded with a high density of mushroom-shaped and stubby spines, which can be stained with fluorescently labeled phalloidin by virtue of its ability binds to actin filaments within the spines (Fig. 1C, Ctrl-neuron). Notably, glioblastoma cells retained viability and invasive potential in the interface conditions, mimicking early tumor infiltration into cortical environment [29].

### 2. Immortal U-87 MG but not serum-free patient-derived NU-757 glioblastoma cell lines induces changes in synaptic morphologies

Relying on the dual-interface co-culture system described in Figure 1, we next sought to determine whether acute exposure to GB cells elicit measurable changes in human cortical synaptic architecture. This platform, which was designed to preserve spatial fidelity and allow controlled, non-overlapping interaction between differentiated human neurons and GB cells, provided a unique opportunity to study early, cell-type-specific effects on synaptic morphology before irreversible structural degeneration occurs.

To assess the potential functional differentiation of the system, we compared the effects of two glioblastoma models: the commonly used immortalized serum-adapted U-87 MG line [30] and the serum-free patient-derived NU-757 line [31], both introduced at the base of the microglia-supported interface. Human iPSC-derived cortical neurons at day 35 of differentiation, featuring mature dendritic arbors and high spine density (Fig. 1B, C, Ctrl-neuron), were exposed to GB cultures over a 24-hour period. Importantly, the use of a human microglia layer in the interface did not compromise the cell viability and neuronal structural integrity, thereby modeling the dynamic and reciprocal signaling present in early tumor infiltration of brain parenchyma.

Neurons co-cultured with U-87 MG cells exhibited pronounced synaptic disorganization. Phalloidin staining showed a loss of mature spine types, particularly mushroom-shaped spines, suggesting cytoskeletal remodeling at excitatory synapses. In contrast, exposure to the serum-free cell line cultured from patient tumor tissue NU-757 did not induce significant alterations in synaptic morphology but an increase of spines compared to control cultures (Fig. 1C, D). On the other hand, Western blot analysis revealed differential expression of synaptic proteins, including presynaptic synaptophysin and postsynaptic PSD-95 and NMDAR2B, between the two glioblastoma models (Fig. S1). If neurons exposed to U-87MG presented only a significant downregulation of PSD-95 proteins, the expression of all three proteins was significantly reduced in those exposed to NU-757 GB cells. These differences likely reflect distinct modes of tumor-neuron interaction rather than differences in intrinsic tumor aggressiveness, highlighting how variant these two GB cells can elicit divergent neuronal responses within the tumor microenvironment.

These findings underscore both the sensitivity and versatility of our co-culture system in capturing subtle effects on neural circuits.

### 3. Acute glioblastoma exposure triggers disease-associated proteomic signatures in healthy human neurons

Given the pronounced synaptic remodeling observed following acute exposure to glioblastoma (GB) cells, we next asked whether these synaptic alterations were accompanied by an acute global shift in protein expression in neuron. Using our dual-interface co-culture model, we performed an unbiased proteomic analysis to capture neuronal responses at the protein level during the critical early window of tumor-neuron interaction.

To isolate neuron-specific proteomic changes, human iPSC-derived cortical neurons were cultured normally (control) or exposed for 24 hours to either U-87 MG or NU-757 GB cells in the dual-interface system (Fig 1A). Following exposure, neuronal cultures were carefully dissociated from the upper layer, excluding GB and microglial layers, and subjected to tandem mass tag (TMT)-based quantitative mass spectrometry (MS). Across conditions, over 5,500 proteins were reliably quantified (Fig. S2).

Neurons exposed to U-87 MG cells exhibited a robust proteomic shift relative to controls, with 520 proteins significantly altered (adjusted p < 0.05, fold change > 1.5).

In sharp contrast, neurons co-cultured with patient-derived NU-757 glioblastoma cells showed a much more constrained proteomic response. Only 83 proteins were differentially expressed compared to controls.

Gene Ontology (GO) and pathway enrichment analyses of the proteomic data from neurons exposed to U-87 and NU-757 glioblastoma cells revealed significant changes in key neuronal structural and functional pathways (Fig. 2A, S3A and B). Both models showed enrichment in terms related to synaptic organization, including presynaptic and postsynaptic compartments, axonal structure, cell junctions, and neuron projection development (Fig. 2A). These pathways collectively point to a disruption in neuronal connectivity and communication, indicating that glioblastoma exposure broadly affects the integrity of neuronal networks. This molecular evidence aligns with our structural observations: neurons exposed to U-87MG cells exhibited reduced MAP2 expression (Fig. S1B) and dendritic spine loss (Fig. 1C), while those exposed to NU-757 retained more typical neuronal architecture. Interestingly, despite these differences in structural responses, both tumor models induced proteomic signatures associated with high-grade and malignant glioma, as revealed by DisGeNET analysis (Fig. 2B). This suggests that the glioblastoma microenvironment, regardless of the type of GB cells, triggers a shared set of disease-relevant molecular changes in neurons, particularly targeting pathways essential for maintaining synaptic integrity and neuronal projection. These findings highlight a converging mechanism by which glioblastoma perturbs neuronal function, with potential implications for both tumor progression and neurological decline in affected patients. Importantly, this finding highlights the capacity of our co-culture system to detect the molecular consequences of tumor exposure in a cell-type-specific and time-sensitive manner.

**Figure 2.**
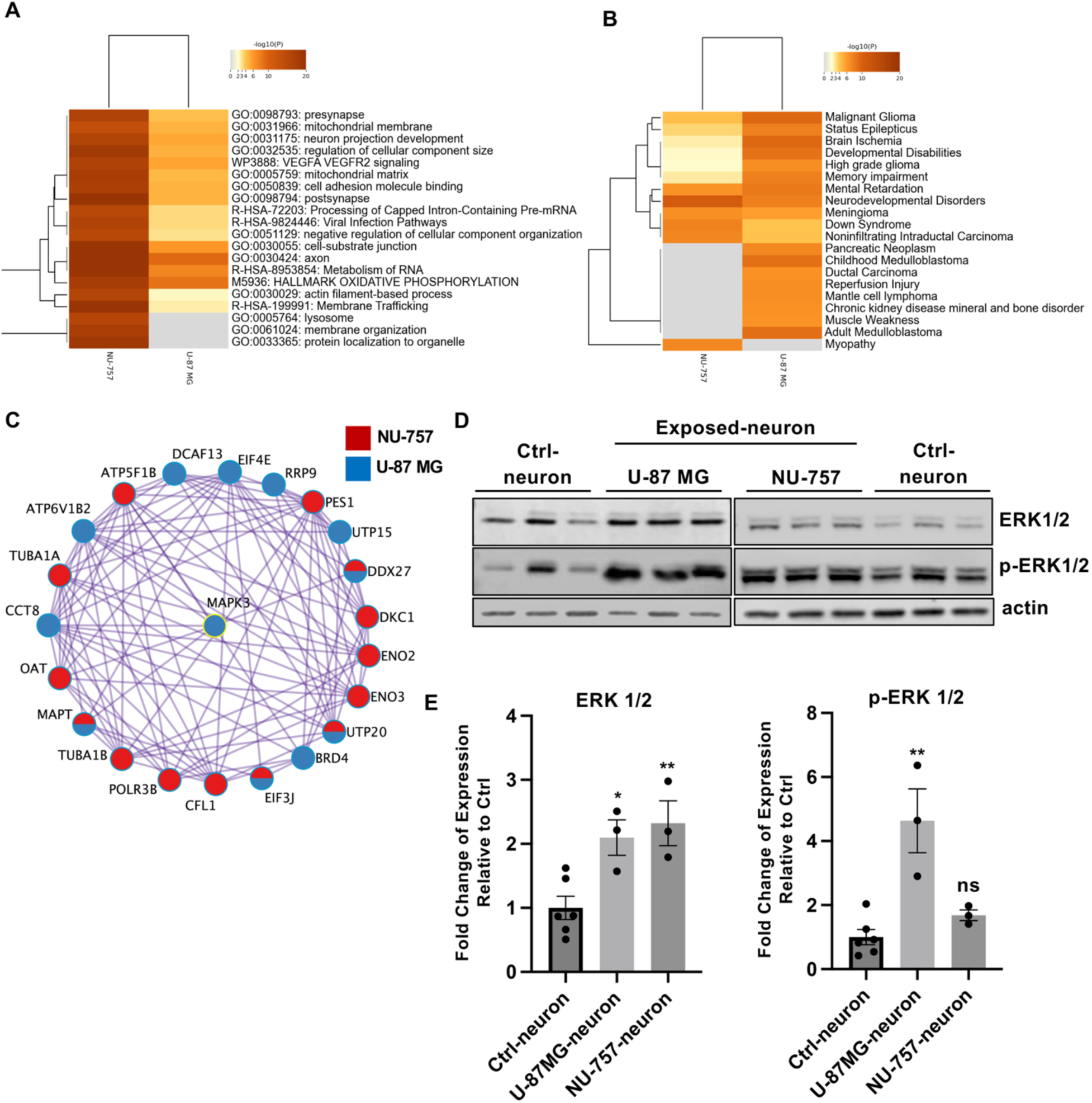
Proteomic analyses reveal distinct signaling pathway activation in neurons exposed to different glioblastoma cells. (A) Heatmap showing pathway and process enrichment analysis of differentially expressed proteins in neurons exposed to NU-757 versus U-87 MG glioblastoma cells for 24 hours. Enrichment was performed using described ontology sources. Heatmap cells are color-coded by p-value; white cells indicate no significant enrichment for that term in the respective protein set. (B) Heatmap of DisGeNET disease enrichment analysis based on differentially expressed neuronal proteins after 24-hour exposure to NU-757 versus U-87 MG glioblastoma cells. Heatmap cells are colored by p-value, with white cells indicating no enrichment. (C) Protein-protein interaction (PPI) network centered on ERK1/2 (MAPK3), highlighting significantly altered proteins in neurons exposed to NU-757 (red) or U-87 MG (blue) glioblastoma. (D) Immunoblot analysis of neuronal lysates from dual-interface co-cultures, including control neurons (Ctrl-neuron), neurons exposed to U-87 MG (U-87 MG-neuron), and neurons exposed to NU-757 (NU-757-neuron). Blots were probed for total ERK1/2, phosphorylated ERK1/2 (p-ERK1/2), and actin as a loading control. Data represent three independent experiments (n = 3). (E) Quantification of ERK1/2 and its phosphorylated form, p-ERK1/2 immunoreactivity in the blots shown in D. Data shown is the mean ± SEM. Statistical analysis was performed using unpaired t-tests; Significance is indicated as: ns (not significant), *P < 0.05, **P < 0.01

In sum, these data suggest that early exposure to GB-derived signals can trigger neuronal reprogramming toward a stress-related, disease-like state, potentially priming the brain microenvironment for long-term dysfunction. The integration of proteomic profiling into our model offers a valuable readout for mechanistic investigation and therapeutic targeting of tumor-neuron communication.

### 4. Glioblastoma induces activation and alters subcellular distribution of MEK/ERK pathways in peritumor neurons

To elucidate the intracellular pathways underpinning glioblastoma-induced synaptic and molecular alterations, we focused on the MEK/ERK cascade, a critical mediator of neuronal stress responses and synaptic remodeling. Building on proteomic pathway enrichment (Fig. 2C), we performed immunoassays on iPSC-derived cortical neurons after 24 hours of GB exposures. Western blot (WB) (Fig. 2D) and immunofluorescence (IF) (Fig. 3A and B) analysis show significant changes in both ERK1/2 total levels and phosphorylation levels of ERK1/2 (p-ERK1/2) in GB-exposed to neurons. High-resolution immunofluorescence to localize these proteins in neurons show that U-87 MG exposure exhibited significantly elevated p-ERK levels in different neuron compartments, in the nucleus and soma but not dendrites, with no change (p=0.7, sample size: 1102-2186 locations) in spine-localized p-ERK puncta (Fig. 3C, E) and ∼4.2-fold decreases (p<0.0001, sample size: 81-111 locations) in nuclear/cytoplasm ratio of p-ERK intensity (Fig. 3C, E) compared to controls. In contrast, neurons exposed to patient-derived NU-757 cells showed no significant changes in p-ERK nuclear/cytoplasm ratio (p=0.06, sample size: 72-111 locations) (Fig 3C, D), but ∼4 times increases (p<0.0001, sample sizes: 1349-2186 locations) at spines location (Fig. 3C, E), consistent with their preserved synaptic morphology distinct from U-87 MG (Fig 1C).

**Figure 3.**
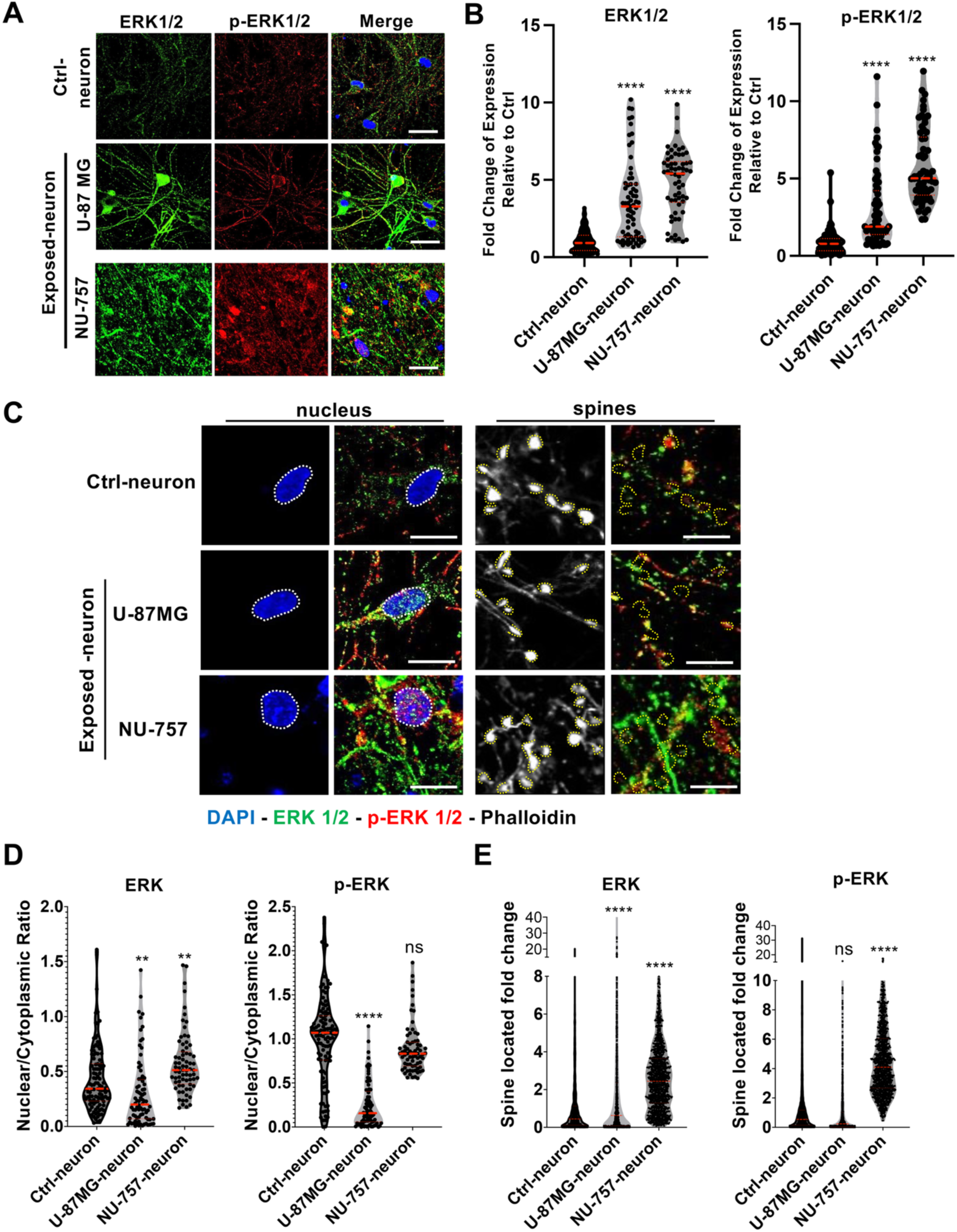
Differential subcellular localization and alteration of ERK1/2 signaling in neurons exposed to glioblastoma cells. (A) Representative immunofluorescence images showing total ERK1/2 (green) and phosphorylated ERK1/2 (p-ERK1/2, red) in neurons exposed for 24 hours to U-87 MG or NU-757 glioblastoma cells, compared to controls. Nuclei were stained with DAPI (blue). Scale bars = 50 µm. (B) Quantification of ERK1/2 immunofluorescence intensity in neuronal cultures exposed to U-87 MG or NU-757 glioblastoma for 24 hours versus controls. Data represent pooled measurements from 70-120 randomly selected neuronal fields per independent experiment. Values are mean ± SD. Statistical analysis was performed using unpaired t-tests; significance: ****P < 0.0001. (C) Representative images of triple immunofluorescence staining for phalloidin (gray), total ERK1/2 (green), and p-ERK1/2 (red) highlighting different neuronal compartments after 24-hour glioblastoma exposure. Dotted lines outline nuclei (DAPI, blue) and intact dendritic spines as indicated by phalloidin staining. Scale bars = 20 µm and 5 µm for nucleus panels and spine panels, respectively. **(D-E)** Quantification of ERK1/2 immunofluorescence intensities in distinct neuronal compartments from images in (C). (D) Ratio of ERK1/2 levels in nucleus versus cytoplasm, pooled from 72-111 locations across three independent experiments. (E) ERK1/2 levels at spine locations, based on 1120-2186 measurements from three independent experiments. Data are presented as mean ± SD. Statistical comparisons used unpaired t-tests. Significance is indicated as: ns (not significant), **P < 0.01, ****P < 0.0001.

Expanding this analysis, we assessed additional MAPK family members, mixed lineage kinase 2 (MLK2) and p38 MAPK, both of which are implicated in regulating cytoskeletal dynamics and dendritic spine morphology. MLK2, is significantly upregulated in proteomes of U-87MG-exposed neurons (Fig. S3C, MAPK310), which are known to activate downstream MAPK pathways including JNK and p38, has been shown to modulate actin cytoskeleton remodeling and spine stability, while p38 MAPK regulates synaptic plasticity and spine pruning via phosphorylation of cytoskeletal-associated proteins. WB analysis of GB-exposed neurons shows MLK2 and phospho p38 levels significantly upregulated in U-87 exposure but not in NU-757 exposure compared to control neuron samples (Fig. S4A). Interestingly, the following analysis of subcellular distribution of these proteins in neurons provided another informative perspective. Neurons exposed to NU-757 cells showed a significant increase in nucleus-localized MLK2 level (∼1.3-fold) but not in nuclear p38 level relative to controls (Fig. S4B, C). This elevated nuclear MLK2 level is in contrast with neurons exposed to U-87MG cells, where MLK2 remained at baseline for both nucleus and spine locations (Fig. S4C). At spine locations, neurons exposed to NU-757 cells showed a significant increase for both MLK2 (∼2.0-fold), and p38 (∼1.9-fold). Neurons exposed to U-87MG cells showed a significant increase in p38 (∼1.3-fold). This differential activation correlates with the distinct synaptic phenotypes observed: U-87 MG exposure induces spines loss and aberrant morphology, while NU-757 exposure preserves spine density and structure (Fig. 1C).

The selective upregulation of p38 family in U-87 MG-exposed neurons in both nucleus and spine locations likely contributes to destabilization of actin filaments and dendritic spines, as these kinases are known to phosphorylate cytoskeletal regulators such as MAP2 and PSD95, leading to spine shrinkage and synaptic weakening. Conversely, the absence of this activation in nucleus locations of NU-757 conditions may explain the maintenance of normal spine morphology despite tumor exposure.

Crucially, these findings underscore the unique capacity of our dual-interface co-culture model (Fig. 1A) not only to discriminate glioblastoma cells-specific effects on neurons but also to resolve signaling events with subcellular precision, capturing dynamic and compartmentalized neuronal responses in both acute and potentially chronic tumor microenvironments.

### 5. MEK pathway inhibition potentially reverses spine loss in U-87 MG-exposed neurons and impairs glioblastoma growth and migration

Having established that U-87 MG glioblastoma cells induce MEK/ERK-dependent signaling in neurons exposed to them (Fig. 2 and 3), we next tested whether pharmacological inhibition of this pathway could reverse synaptic disruption and impair GB cells migration/invasion. Human iPSC-derived cortical neurons were treated with the MEK1/2 inhibitor Selumetinib [32], an FDA-approved drug for neurofibromatosis type 1 (NF1), or an ERK inhibition peptide [33], during the 24-hour co-culture period with U-87 MG cells in the dual-interface system. This MEK and ERK inhibitions partially restored neuronal morphology at the dendritic spines level. Phalloidin staining of actin-rich dendritic spines revealed a significant increase of spine density, (Fig. 4B and E). Quantitative imaging confirmed suppression of phosphorylated ERK (p-ERK) in neurons, validating effective MEK pathway blockade (Fig. 4A, C and D).

**Figure 4.**
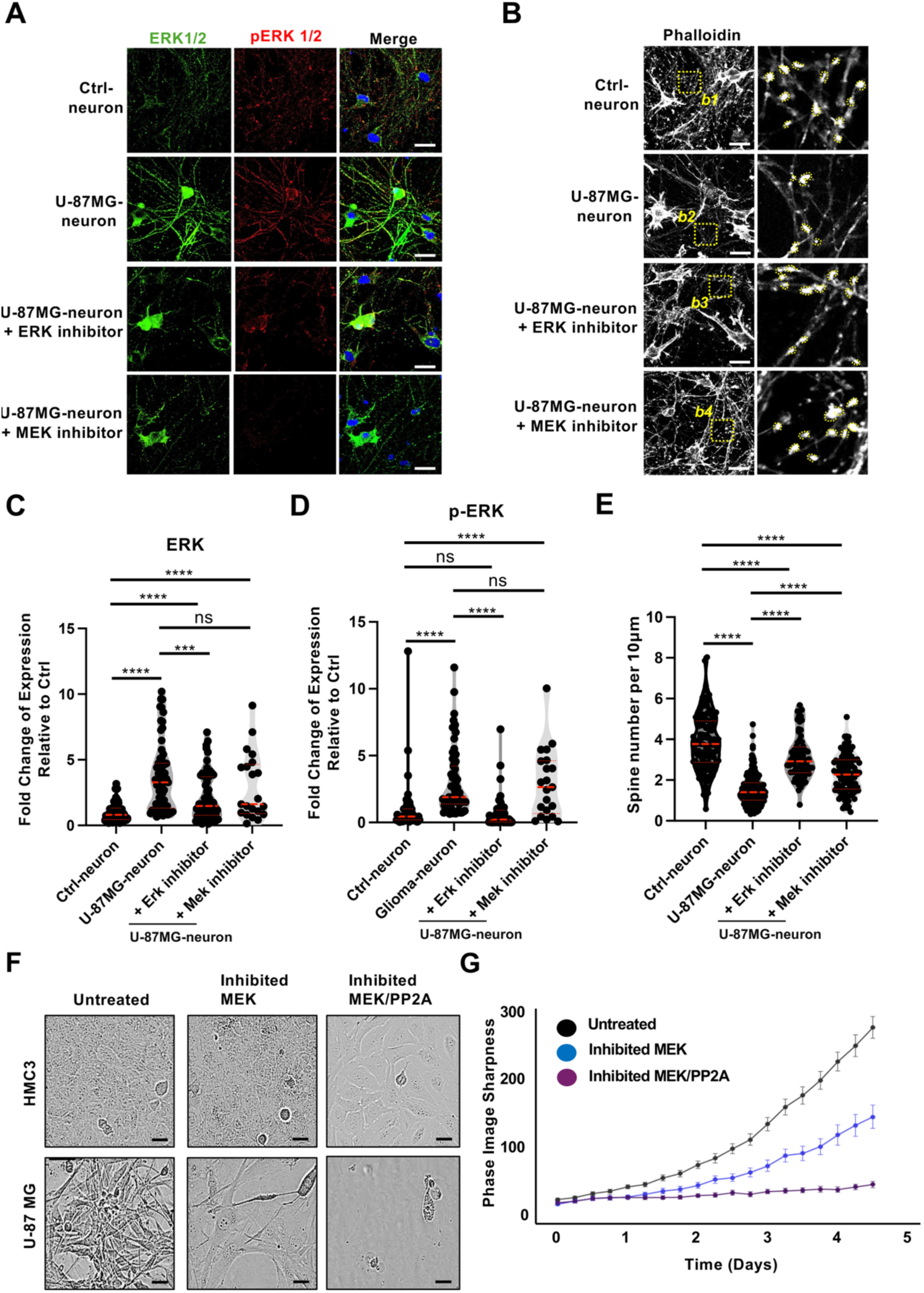
MEK/ERK signaling inhibition reduces glioblastoma-induced neuronal ERK activation and spine loss. **(A)** Representative immunofluorescence images of total ERK1/2 (green) and phosphorylated ERK1/2 (p-ERK1/2, red) in neurons under different conditions: control (Ctrl-neuron), exposed to U-87 MG glioblastoma cells (U-87 MG-neuron), U-87 MG-exposed neurons treated with an ERK peptide inhibitor (U-87 MG-neuron +ERK inhibitor), or U-87 MG-exposed neurons treated with the MEK inhibitor Selumetinib (U-87 MG-neuron +MEK inhibitor). Nuclei were stained with DAPI (blue). Scale bars = 30 µm. **(B)** Fluorescent phalloidin staining (gray) of dendritic spines in neurons after 35 days of differentiation under the same conditions as in (A). Insets (b1-b4) show magnified views of the dashed regions. Scale bars = 30 µm. **(C-D)** Quantification of immunofluorescence intensity of total ERK1/2 (C) and p-ERK1/2 (D) in neurons under each condition from (B). Data are pooled from 21-67 randomly selected neuronal fields per independent experiment. Values represent mean ± SD. Statistical comparisons were performed using unpaired t-tests. Significance: ns (not significant), ***P < 0.001, ****P < 0.0001. (E) Quantification of dendritic spine density (spines per 10 µm) under each condition from (B). Data represent 72-160 dendritic segments from 2-3 independent experiments. Values are shown as mean ± SD. Unpaired t-tests were used for statistical analysis; ****P < 0.0001. **(F-G)** Phase-contrast images (F) and quantification (G) of HMC3 microglia and U-87 MG cells imaged using Incucyte. Treatment with Selumetinib (MEK inhibitor) or Selumetinib combined with DT-061 (MEK/PP2A pathway inhibition) reduced U-87 MG cell proliferation and migration, and induced cell death. Scale bars = 30 µm. Values are shown as mean ± SEM. Data are pooled from 4 cultures per each condition; 5 fields were imaged per each culture.

In the same conditions, U-87 MG glioblastoma cells exhibited significantly impaired expansion and migration. MEK inhibition by Selumetinib reduced the two-dimensional surface coverage of U-87 MG cells by ∼55% (p < 0.001) and blocked their transwell invasiveness (Fig. 4F, G and Fig. S5C). The viability of the treated-cells was not affected, suggesting that MEK inhibition alone does not induce cytotoxicity.

To identify complementary targets that might amplify the anti-tumor effect, we examined our proteomic data (Section 3, Fig S5A) for neuronally regulated factors with tumor-suppressive potential. Notably, Protein Phosphatase 2A (PP2A) regulatory subunits were downregulated in neurons exposed to U-87 MG (Fig. S5B), suggesting a glioblastoma-driven suppression of this critical phosphatase. Given PP2A’s known role in antagonizing oncogenic MAPK signaling and stabilizing cytoskeletal architecture, we hypothesized that PP2A activation could synergize with MEK inhibition.

Treatment with the small-molecule PP2A activator DT-061 alone showed ineffective in U-87 MG cell proliferation and migration, except 10 µM concentration of treatment (Fig S6A). However, a combination of DT-061 and Selumetinib suppressed GB cells expansion and trans-well invasion, and notably, induced cytotoxic death in U-87 MG cells compared with the mock and single treatments (Fig. 4F, G and Fig. S5 E, F). This synergistic effect likely reflects a collapse of compensatory kinase-phosphatase balance within GB cells and highlights a promising therapeutic interaction between phosphatase reactivation and kinase inhibition.

Importantly, this dual-target approach had no toxic effects on control neurons or neurons co-cultured with patient-derived NU-757 glioblastoma cells compared to serum-adapted GB U-87cells. Moreover, NU-757 cells showed no significant change in growth, migration, or viability under MEK inhibition (Fig. S5D). This differential response suggests that NU-757 cells rely less on MAPK signaling, consistence with NU-757-exposed neuronal proteomes data, and instead depend on distinct molecular pathways for proliferation and survival. On the other hand, the inhibitors were evaluated for their toxicity on normal neurons (Fig. S6E). These findings underscore the biological heterogeneity of GB, and the importance of tailored therapeutic strategies based on tumor type and signaling context, a point we further explore in the Discussion.

## DISCUSSION

The interaction between GB and neurons is increasingly recognized as a bidirectional process that not only shapes the progression of the tumor but also contributes to neurological dysfunction in patients [34]. Furthermore, current GB treatments fail to address the invasive aspect of tumor spread, which is responsible for recurrence and disease progression. Thus, there is an urgent need for a better understanding of tumor biology at the infiltrative margin, as well as the development of improved combination therapies that address not only tumor proliferation but also migration for all GB patients. In this study, we established a novel dual-interface co-culture system to model early, spatially controlled interactions between human GB cells and iPSC-derived cortical neurons. This system allowed us to uncover tumor cell-specific effects on neuronal morphology, proteomic reprogramming, and intracellular signaling with subcellular resolution, and to identify actionable therapeutic vulnerabilities that simultaneously target tumor and neuronal compartments.

One of the key innovations of this platform lies in its capacity to model acute tumor-neuron interactions with defined spatial architecture, overcoming the limitations of static organoids and mixed co-cultures. The system facilitates the separation of neurons from GB cells following exposure, enabling a precise investigation of neuron-specific pathogenic events that precede GB cells proliferation and invasion. The use of mature, synaptically active human cortical neurons layered above a microglial monolayer provides a biologically relevant context in which GB-secreted factors can influence neurons without physical overgrowth. This design enables the resolution of early molecular and functional changes that precede irreversible structural damage, offering a tractable window for intervention and mechanistic dissection.

Using this system, we discovered that the commonly used immortalized glioblastoma line U-87 MG, but not the serum free patient-derived NU-757 glioblastoma cells, induced rapid and significant remodeling of neuronal dendritic spines. Neurons exposed to U-87 MG cells displayed reductions in spine density, alterations in spine morphology, and disruptions in synaptic protein distribution. These structural and synaptic modifications occur before the functional integration of GB cells into the neuron’s circuits and before neuronal dysfunction. This finding is supported by Svenja K. Tetzlaff et al. [35]. They noted that in certain patient-derived models, the majority of GB cells and neurons form early functional connections during GB invasion, prior to the onset of neuronal dysfunction and hyperexcitability, which appear later in the disease [35]. The changes observed in our study were accompanied by a robust proteomic shift toward degenerative, inflammatory, and cytoskeletal remodeling pathways, while neurons co-cultured with NU-757 cells maintained synaptic structure. This divergence underscores the importance of glioblastoma cell of origine in shaping neuronal responses and highlights the utility of our system for modeling such subtle differences. In addition, the differential outcomes between U-87 MG and NU-757 highlight the limitations of using immortalized serum adapted lines as generalizable GB models and emphasize the need for incorporating serum free patient-derived cells that reflect the phenotype of the tumor type of origin. It is also well documented that GB cell lines including U-87MG present discrepancies in their mutational profiles and their original tumor tissue suggesting that they have deviated from their initial characteristics after a certain amount of time in culture [36]. This could also explain the differences observed.

Our data show that neurons exposed to U-87MG, in addition to the loss of dendritic spines (Fig. 1C), exhibit reduced MAP2 expression (Fig. S1B). MAP2 is a neuronal protein that works as dendrites and dendritic spine marker, interacting with microtubules to maintain dendritic shape [37] and its downregulation indicate structural degeneration and impaired synaptic integrity. While previous studies have highlighted the ability of GB cells to form synaptic connections with neurons, enhancing tumor growth through neuron-to-glioma glutamatergic signaling [38, 39], our findings suggest that this U-87 MG neuronal interaction may come at the expense of normal neuron-neuron connectivity. The decrease in dendritic spines suggests that neuron-glioma interactions serve a dual purpose: they facilitate tumor progression while simultaneously impairing neuronal health. This highlights the complex and detrimental impact of GB on surrounding neural networks.

Mechanistically, we identified activation of the MEK/ERK signaling cascade as a central mediator of the neuronal response to U-87 MG glioblastoma cells. High-resolution imaging revealed ERK activation not only in the neuronal soma and nucleus but also at dendritic spines, suggesting concurrent modulation of transcriptional programs and local synaptic plasticity. This response was further supported by selective changes of MLK2 and p38 MAPK, two additional stress-responsive kinases with known roles in cytoskeletal regulation and synaptic destabilization [40]. Notably, these pathways were not activated similarly in neurons exposed to NU-757 cells. The absence of comparable pathway activation in neurons exposed to NU-757 cells potentially explains the mechanistic underpinnings of the observed differences in neuronal morphology.

From a therapeutic perspective, our data demonstrate that pharmacological MEK inhibition restores dendritic spine morphology and synaptic protein expression in U-87 MG-exposed neurons. While MEK inhibition alone attenuated tumor cell spreading and invasion, it did not induce tumor cell death. However, by integrating proteomic findings from the neuronal compartment, we identified induced expression of PP2A components as a potential tumor-promoting mechanism. Co-treatment with a PP2A activator (DT-061) and MEK inhibitor not only fully suppressed U-87 MG expansion but also induced significant tumor cell death. This combinatorial approach highlights a rational strategy to co-target kinase and phosphatase imbalances that drive tumor aggressiveness and neuronal dysfunction.

Importantly, NU-757 cells were resistant to MEK inhibition, showing no impairment in growth or invasion. Again, this difference likely reflects intrinsic signaling diversity among glioblastomas. While U-87 MG appears to be dependent on MAPK signaling, NU-757 may rely on alternative oncogenic drivers that are not targeted by MEK inhibitors. These findings emphasize the need for personalized therapeutic strategies and caution against generalizing results from immortalized glioma lines to patient-derived models. Our system provides a robust platform for identifying such differential dependencies and testing tailored interventions.

In summary, this study introduces a versatile, compartmentalized in vitro model that enables mechanistic and therapeutic exploration of glioblastoma-neuron interactions at unprecedented resolution. Using this platform, we demonstrate that tumor subtype shapes neuronal vulnerability, identify MEK/ERK and PP2A as convergent signaling nodes at the tumor-neuron interface, and validate a dual-compartment therapeutic approach that restores neuronal structure while suppressing glioblastoma growth. This work advances our understanding of neuro-glioma crosstalk and offers a powerful tool for preclinical modeling of therapeutic strategies that address both tumor burden and neural integrity in glioblastoma.

A limitation of this study is the lack of simultaneous secretome analysis from neurons and GB cells, which prevents a precise mechanistic description of the soluble signaling components that drive their functional connections. Furthermore, we used a specific neuronal subtype, limiting the generalizability. GB tumors form in various brain regions innervated by different neuronal populations, suggesting that the observed relationships are most likely dependent on specific local microenvironments. Therefore, future studies should investigate reciprocal secretome profiles to identify the molecular mediators that underpin neuron-GB communication and systematically evaluate how different neuronal subpopulations, particularly cholinergic neurons, given their crucial neuromodulatory role influence these events. Additionally, tumor chunk cultures, which retain the tumor microenvironment (TME) of their patient of origin, could represent an optimal model for delineating neuron-specific responses to individual tumors.

## MATERIALS AND METHODS

### Cell line cultures

The serum adapted U-87 MG (ATCC HTB-14) and HMC3 (ATCC CRL-3304) cell lines were maintained in Dulbecco’s Modified Eagle Medium (DMEM; Gibco, Cat. #1196511) supplemented with 10% fetal bovine serum (FBS; Gibco, Cat. #1600044), 100 U/mL penicillin, and 100 μg/mL streptomycin (Gibco, Cat. #15070063). Cultures were incubated at 37 °C in a humidified atmosphere containing 5% CO₂.

The serum free patient-derived glioblastoma cell line NU-757 originated from primary human brain tumor tissue, obtained as de-identified specimens from the Northwestern University Nervous System Tumor Bank[41, 42]. NU-757 cells were cultured under adherent, serum-free conditions in phenol red–free Neurobasal medium (Gibco, Cat. #12348017) supplemented with B-27 Supplement minus Vitamin A (Gibco, Cat. #12587010), human FGF-2 (10 ng/mL; Miltenyi Biotec, Cat. #130-097-751), human EGF (10 ng/mL; Miltenyi Biotec, Cat. #130-093-842), GlutaMAX (1×; Gibco, Cat. #35050061), sodium pyruvate (1 mM; Gibco, Cat. #11360070), and penicillin/streptomycin (100 U/mL and 100 μg/mL, respectively; Gibco, Cat. #15070063). NU-757 cells were seeded onto tissue culture plates pre-coated with Geltrex LDEV-free stem cell, a qualified reduced-growth factor basement membrane matrix (Life Technologies, Cat. #A1413302). Coating was performed by incubating plates with Geltrex for 4 hours at 37 °C or overnight at 4 °C. For passaging, NU-757 cells were dissociated into single-cell suspensions using TrypLE Express (phenol red–free; Gibco, Cat. #12604013).

### Human iPSCs and iPSC-derived cortical neuron culture

Human induced pluripotent stem cells (iPSCs) of the KOLF2.1 line (Jackson Laboratory for Genomic Medicine) were cultured in mTeSR™ Plus medium (STEMCELL Technologies) on Matrigel-coated 6-well plates (BD Matrigel™ hESC-qualified Matrix, Cat. #354277; coating time for 30 minutes at room temperature). Routine passaging was performed using ReLeSR™ (STEMCELL Technologies), an enzyme-free dissociation reagent, and medium was replaced daily to support optimal cell growth.

Differentiation of iPSCs into cortical neurons was carried out as previously described [27], using an established neurogenin-2 (NGN2)-induction protocol [26, 43]. Briefly, human iPSCs stably carrying the NGN2 transgene were dissociated into single cells and plated on Matrigel-coated plates in Induction Medium (IM) supplemented with doxycycline (2 µg/mL) to induce NGN2 expression and the ROCK inhibitor Y-27632 (10 µM; Tocris Bioscience, Cat. #1254) to promote cell survival. Cells were maintained in induction conditions for 3 days. On Day 4 of differentiation, cells were detached using Accutase (Gibco, Cat. #A1110501) and replated at defined densities onto poly-D-lysine (PDL; Gibco, Cat. #A3890401) and poly-L-ornithine (PLO; Sigma, Cat. #P4957-50mL)-coated 12-well plates. For coating, each well received 700 µL of PDL solution, which was incubated for 4 hours at room temperature. PDL was then removed and replaced with an equal volume of PLO solution, which was incubated overnight at 4 °C. Before cell seeding, the wells were rinsed three times with sterile PBS (5 minutes per wash).

Following replating, cells were cultured in Neuronal Culture Medium (CM). Medium was partially replaced (50%) daily using pre-warmed CM. Neurons were maintained for at least 35 days prior to experimental use. Day 35 neurons were used for all downstream analyses.

### Dual-interface co-culture system

This interface human neuronal culture system is adapted from previously described methods for culturing rodent embryonic hippocampal neurons on coverslips suspended above a glial monolayer [28, 44, 45]. Briefly, glass coverslips were first sterilized by autoclaving. Once dry, three wax legs were applied to each coverslip to act as spacers. To create the legs, a Pasteur pipette was dipped into molten paraffin and used to place three evenly spaced dots near the periphery of each coverslip. The wax legs were approximately 0.5 mm in height and 1.0-2.0 mm in diameter, providing physical separation between the neuronal and glial layers during co-culture.

After the wax solidified, each coverslip was transferred to a well of a 12-well culture plate and sterilized under UV light for one hour. Coverslips were then coated sequentially with poly-D-lysine and poly-L-ornithine as described above to promote cell adhesion. Human induced pluripotent stem cells (iPSCs) were subsequently seeded and differentiated directly on the coated coverslips into mature cortical neurons as described above.

For co-culture experiments, differentiated neurons on coverslips were suspended above a monolayer of human microglial HMC3 cells seeded with either U-87 MG or serum free patient-derived NU-757 glioblastoma cells. Neurons were maintained in this dual-interface configuration for 24 hours prior to downstream analyses.

### Neurons and glioblastoma co-culture in the dual-interface culture system

In this system, GB and microglial cells were first co-seeded onto 12-well plates one day prior to the addition of neurons (Day 1). Cells were plated at a 1:1 ratio, while control wells contained only HMC3 cells. Plates were incubated overnight at 37 °C with 5% CO₂. On the day of co-culture initiation (Day 0), the medium in the GB/microglia cultures was replaced with fresh neuronal culture medium (CM). Coverslips with differentiated neurons were then carefully lifted from their original plates and inverted onto the top of the GB/HMC3 or control HMC3 cell layers. This orientation positioned the neuronal layer facing the GB/microglia layer while maintaining a defined physical separation of approximately 0.5 mm, created by the wax feet on the coverslips.

This dual-interface setup enabled exchange of diffusible factors between neurons and GB/microglia cells without allowing direct cell-cell contact. Co-cultures were incubated for 24 hours before neurons were collected for downstream analyses, including proteomics, western blotting, or immunostaining.

### Western blot analysis

Cells were collected and lysed in 1× cell lysis buffer (Cell Signaling Technology, Cat. #9803) supplemented with a protease and phosphatase inhibitor cocktail (Thermo Scientific, Cat. #A32959). Total protein concentrations were determined using the bicinchoninic acid (BCA) assay (Pierce, Cat. #23225), and samples were stored at-80 °C until further use.

For electrophoresis, 10-25 μg of total protein per sample were resolved on 4-20% gradient Bis-Tris precast gels (GenScript, NJ, USA) and transferred to 0.45 μm nitrocellulose membranes (Bio-Rad, CA, USA). Membranes were blocked with 5% bovine serum albumin (BSA; Gold Biotechnology, Cat. #A-420-500) in TBS buffer for 1 hour at room temperature, then incubated overnight at 4 °C with the primary antibodies (listed in Table S1). Following primary antibody incubation, membranes were washed and incubated with secondary antibodies (listed in Table S1). Protein bands were visualized using the LI-COR Odyssey Imaging System. Densitometric quantification of Western blot bands was carried out using ImageJ software.

### Immunocytochemistry

Cells grown on glass coverslips were washed with phosphate-buffered saline (PBS, 1x) and fixed in 4% paraformaldehyde for 12 minutes at room temperature. For detection of intracellular antigens, cells were permeabilized with 0.1% Triton X-100 in PBS for 5 minutes prior to blocking. Non-specific binding was blocked with 1% bovine serum albumin (BSA) in PBS for 30 minutes at room temperature (RT).

Primary antibodies (listed in Table S1) were diluted in 1% BSA/PBS and applied to cells for 2 hours at RT. After washing with PBS, fluorophore-conjugated secondary antibodies diluted in 1% BSA/PBS were applied for 1 hour at room temperature in the dark. Cells were then washed five times (5 minutes each) in PBS and counterstained with DAPI (Invitrogen, Cat. #D1306) to visualize nuclei. Coverslips were mounted using Vectashield Mounting Medium (Vector Laboratories, Cat. #H-1900) and stored at 4°C until imaging.

All imaging was performed using Olympus FV3000 Multi-Photon Confocal Microscopes with identical acquisition settings across conditions. A 63x oil immersion objective (numerical aperture, NA=1.4) was used for high-resolution image capture.

### Dendritic Spine Analysis

Dendritic spine density was assessed to evaluate synaptic effects, as previously described [27, 44]. Neurons were exposed in different culture conditions, followed by fixation in 4% paraformaldehyde. Neurons cultured on coverslips were then stained with either Alexa Fluor 488-phalloidin or rhodamine-phalloidin to label F-actin-rich dendritic spines. Confocal images were acquired using Olympus FV3000 Multi-Photon Confocal Microscopes equipped with 63x oil and 100x oil immersion objectives (NA = 1.4 numerical aperture, NA=1.4).

For quantification, 5-10 isolated dendritic segments were selected per image. Images were processed in ImageJ (FIJI) using a threshold optimized to include dendritic spine outlines while excluding background fluorescence [46]. Spine numbers were normalized to the corresponding dendritic length and reported as spines per micrometer (spines/μm). For each condition, 5-10 images were analyzed across 2-3 independent cultures.

### Immunofluorescence Intensity Quantification

Fluorescence intensity within defined regions of interest (ROIs) was quantified using ImageJ (FIJI). For each sample, ROIs were manually selected based on the area of interest, such as dendritic spines, dendrites, or intracellular compartments. Each image was processed to ensure uniformity, including background subtraction and optimization of thresholding to exclude non-specific fluorescence signals. The mean fluorescence intensity per pixel was calculated by dividing the integrated density by the ROI area. To control for variability between experimental conditions, all fluorescence intensities were normalized to the average intensity of the control group. Data from at least 5 randomly selected fields per condition were quantified to ensure robust sampling. For each condition, 15-24 neurons from 2-3 independent cultures were analyzed.

### Proteomics analysis

#### Sample preparation and protein digestion via STrap

For each proteomics experiment, neuron control samples (neurons cultured in the dual-interface system without glioblastoma cells) were prepared using the same neuronal differentiation batch as those exposed to GB cells (U-87 MG or NU-757) to avoid batch effects. Each culture condition included four biological replicates. Neuron cultures were harvested in a lysis buffer containing 5% SDS in 50 mM triethylammonium bicarbonate (TEAB; Sigma, Cat. #T7408) for protein extraction. Samples were centrifuged at 10,000 × g for 10 minutes at 4 °C, and the resulting supernatant was collected for subsequent proteomic analysis. Total protein concentrations were quantified using a bicinchoninic acid (BCA) protein assay kit (Pierce) according to the manufacturer’s instructions.

Samples were then prepared for trypsin digestion using the S-Trap method. Five microliters of 50 mM ABC containing 5 µg/µL DTT were added to each sample, followed by incubation at 65 °C for 15 minutes. Next, 5 µL of 50 mM ABC containing 15 µg/µL iodoacetamide were added and incubated at room temperature for 15 minutes in the dark. Samples were acidified by adding 12% phosphoric acid (1:10 v/v acid to sample). For every 25 µL of sample, 165 µL of TEAB (1 M)/methanol (10:90 v/v) was added and loaded onto the S-Trap column for further processing.

Samples were centrifuged at 4,000 × g for 3 minutes at 4 °C to remove the supernatant. The column was washed 3-6 times with 150 µL of TEAB/methanol (10:90 v/v), depending on the initial loading volume. After the final wash, sequencing-grade trypsin dissolved in 50 mM TEAB was added, and digestion was carried out overnight at 37 °C.

The next day, peptides were sequentially eluted with 40 µL of 50 mM TEAB, 0.1% formic acid (FA), and 0.1% FA in 50% acetonitrile. The eluates were pooled, dried using a vacuum concentrator, and resuspended in 20 µL of 50 mM acetic acid. Peptide concentration was determined by absorbance at 280 nm using a Nanodrop spectrophotometer.

#### LC-MS/MS on Eclipse

Nano-liquid chromatography-nanospray tandem mass spectrometry (Nano-LC/MS/MS) for protein identification was performed using a Thermo Scientific Orbitrap Eclipse mass spectrometer. Samples (1 µg) were separated on an EASY-Spray nano column (PepMap™ RSLC, C18, 3 µm, 100 Å, 75 µm × 250 mm; Thermo Scientific) using a 2D RSLC HPLC system (Thermo Scientific). Each sample was injected into a µ-Precolumn Cartridge (Thermo Scientific) and desalted with 0.1% formic acid in water for 5 minutes. The injector port was then switched to inject mode, and peptides were eluted from the trap onto the analytical column. Mobile phase A consisted of 0.1% formic acid in water, and mobile phase B was acetonitrile with 0.1% formic acid. The flow rate was set at 300 nL/min. Mobile phase B was increased from 2% to 16% over 105 minutes, then from 16% to 25% over 10 minutes, followed by an increase from 25% to 85% over 1 minute. The column was held at 95% B for 4 minutes before being returned to 2% B in 1 minute. The column was equilibrated at 2% B (98% A) for 15 minutes before the next sample injection.

MS/MS data were acquired with a spray voltage of 1.95 kV and a capillary temperature of 305 °C. The scan sequence was based on the preview mode data-dependent TopSpeed™ method. Full MS scans were recorded between m/z 375-1500, followed by MS/MS scans of the most abundant peaks in the next 3 × 1 seconds. Full scans were acquired in Fourier Transform (FT) mode at a resolution of 120,000 with internal mass calibration for high mass accuracy. Three compensation voltages (CV = - 40,-60, and-80 V) were applied for acquisition. The AGC target for FT full scans was set to 4 × 10⁵ ions, with the maximum injection time set to “Auto” and one microscan. MSⁿ was performed using higher-energy collisional dissociation (HCD) in Orbitrap mode to ensure accurate mass detection of post-translational modifications. The HCD collision energy was set at 30%. The AGC target for ion trap MSⁿ scans was 1 × 10⁴ ions, with an “Auto” maximum injection time and one microscan. Dynamic exclusion was enabled with a repeat count of 1 within 20 seconds and a low and high mass width of ±10 ppm.

#### Database search

The resulting MGF files generated from the samples were searched using Mascot Daemon (Matrix Science, version 2.7.0, Boston, MA) against both the RSSB sequence and the human protein database. The precursor ion mass accuracy was set to 10 ppm, and the search parameters included allowance for the accidental selection of one ^13C peak. Fragment mass tolerance was set to 0.5 Da. Variable modifications considered during the search included acetylation (K), phosphorylation (S and T), oxidation (M), deamidation (N and Q), and carbamidomethylation (C). Up to four missed cleavages by the enzyme were permitted. A decoy database was also searched to determine the false discovery rate (FDR), and peptides were filtered accordingly. The significance threshold was set at p < 0.05, and valid peptide identifications required bold red peptide matches. Proteins were considered valid if they had an FDR of less than 1% and a minimum of two significant peptides. All modified peptides were manually validated.

#### Quantitation and bioinformatics

Relative quantitation was performed using a label-free quantitation approach based on mass spectral peak intensities. Peptide precursor (MS1) intensities from both modified and unmodified peptides were extracted from the ProteomeDiscoverer MASCOT search results and summed for quantitative comparison. Protein intensities were normalized to total protein intensity, and “low abundance resampling” was applied as the imputation mode. Statistical significance of expression differences between groups was evaluated using a student’s t-test. Proteins with a p-value < 0.05 and a fold change ≥ 2 were considered upregulated, while those with a fold change ≤ 0.5 were considered downregulated.

The data underwent bioinformatics analyses using publicly available online databases including, Metascape [47] and Cytoscape [48]. For comparative pathway and network enrichment analysis, protein list enrichment and network analyses were performed using the Metascape platform [47]. Briefly, for each given gene set encoded from different expressed protein list, enrichment analyses have been carried out. Two defined gene sets (“NU-757” and “U-87 MG”) were analyzed independently and in combination. Gene identifiers were mapped to *Homo sapiens* Entrez Gene IDs, and redundancy was resolved prior to analysis. Functional enrichment analyses were conducted across multiple ontology sources including GO Biological Processes, GO Molecular Functions, GO Cellular Components, Reactome, KEGG, Hallmark, and BioCarta. Enriched terms were identified using a cumulative hypergeometric distribution with a cutoff of *p* < 0.01, enrichment factor > 1.5, and a minimum gene count of 3. Multiple testing correction was applied using the Benjamini-Hochberg procedure to calculate *q*-values. To reduce redundancy and visualize the biological landscape, enriched terms were hierarchically clustered using Kappa-statistic-based similarity (>0.3), with the most statistically significant term within each cluster serving as its representative. Up to 20 top clusters were visualized via heatmaps and enrichment networks. Protein-protein interaction (PPI) networks were built using data from STRING (physical score > 0.132), BioGRID, OmniPath, and InWeb_IM. The MCODE algorithm was applied to detect densely connected network modules, and enrichment analysis was performed on each module individually to annotate biological functions. Further characterization was conducted using curated gene set libraries including DisGeNET. All enrichment results were filtered based on standard thresholds (*p* < 0.01, enrichment factor > 1.5) and summarized by most significant terms per cluster. Data visualizations, including network maps (rendered with Cytoscape), were generated to explore shared and unique pathway associations across gene lists.

### Glioblastoma Cell Migration and Invasion Assay

To assess the migratory and invasive properties of glioblastoma (GB) cells, NU-757 and U-87 MG cells were seeded at low density in Boyden chambers with or without Geltrex-coated membranes. For this experiment, Millicell 12-well hanging cell culture inserts (Merck Millipore, cat. PTEP12H48) and HTS Transwell-96 permeable supports (Corning, cat. #3374) with 8.0 µm pore polyester membrane were utilized. The pore size was selected based on the migratory cells being used and the inserts are appropriate for evaluating cell migration and invasion.

For the cell migration/invasion assay, inserts were placed into receiver plates. Pre-warmed cell culture medium without cells was added to the basolateral compartment of each well. Next, cell suspension was prepared and seeded onto the apical side of the inserts: 100 μL per well for 96-well inserts (HTS Transwell-96) and 700 μL per insert for 12-well inserts (Millicell). An identical cell number was seeded across all conditions. Plates were then incubated at 37°C in a humidified CO₂ incubator. Finally, the migrated/invaded cells accumulating in the receiver wells were monitored over time using the IncuCyte system (IncuCyte S3, Sartorius). To validate the assay system and minimize bias, each condition was performed in at least triplicate. Furthermore, a non-invasive cell line, the HMC3 cells, was included as a control to confirm the barrier membrane effectively inhibited invasion.

In inhibition experiments, cells were seed as above and treated twenty-four hours after with varying concentrations of a MEK inhibitor (selumetinib, MedChemExpress, Cat. # HY-50706), ERK inhibitor peptide [33], or a PP2A activator (DT-061, MedChemExpress, Cat. # HY-112929). Plates were placed into the IncuCyte system (IncuCyte S3, Sartorius), housed within a normal tissue culture incubator (37°C, 5% CO₂). The Incucyte® Live-Cell Analysis Systems scan the tissue culture plates at predetermined intervals to monitor cultures and generate quantified kinetic data. Here 5 or 49 images per well were acquired using a 10x objective every six hours over a period of five to seven days. This live-cell imaging allowed for continuous monitoring of migrated cells and cell proliferation and the generation of growth and growth inhibition curves via the basic analyzer function and the phase image sharpness metric, which is automatically calculated by the IncuCyte software. As cells proliferate, they generate additional edges and density, increasing contrast and sharpness. For the analysis, we chose the desired time periods, experimental settings, number of images and the replicates, then, a masking process was initiated to eliminate the background so that the cells underneath could be seen clearly. To have a consistent comparison between the conditions, the starting point was also normalized. Finally, the graph and the raw data generated by the software were exported and used accordingly

## Statistical Analysis

All statistical analyses were conducted using GraphPad Prism version 10.0.0. Data are presented as mean ± standard error of the mean (SEM). Comparisons between two groups were performed using unpaired two-tailed t-tests, while comparisons across multiple groups were assessed using one-way ANOVA followed by Tukey’s post-hoc test for multiple comparisons. Details regarding the number of replicates (n), type of statistical test used, and significance levels are provided in the corresponding figure legends.

## AUTHOR CONTRIBUTIONS

Conceived and designed the experiments: NL NC ON PG MV. Performed the experiments: ON NL NA BF LK LZ. Analyzed the data: ON NA AM MV LZ BF PG NC NL. Wrote the paper: ON NC NL MV.

## ACKNOWLEDGMENTS

We thank Dr. Michael Ward at National Institute of Neurological Disorders and Stroke (NINDS/NIH) for the K4-PB-TO-hNGN2 and K13-EF1a vectors. We thank Dr. William C. Skarnes at The Jackson Laboratory for Genomic Medicine for the human IPSCs lines. We acknowledge recourses from the Campus Microscopy and Imaging Facility (CMIF) and The OSU Comprehensive Cancer Center (OSUCCC) Microcopy Shared Resource (MSR), The Ohio State University (RRID:SCR_025078). This facility is supported in part by grant P30 CA016058, National Cancer Institute, Bethesda, MD. NL was supported by a Warren Alpert Distinguished Scholar Award. NC was supported by NIH grants CA241105 and GM151124. NL, NC and PG were supported by The Daniel J. Zwayer Brain Cancer Research Fund.

## REFERENCES

1. Ostrom, Q.T., et al., CBTRUS Sta*s*cal Report: Primary Brain and Other Central Nervous System Tumors Diagnosed in the United States in 2013-2017. Neuro-Oncology, 2020. 22: p. 1–42.

2. Ostrom, Q.T., et al., CBTRUS Sta*s*cal Report: Primary Brain and Other Central Nervous System Tumors Diagnosed in the United States in 2014-2018. Neuro-Oncology, 2021. 23: p. 1–105.

3. Faisal, S.M., et al., Current Landscape of Preclinical Models for Pediatric Gliomas: Clinical Implica*ons and Future Direc*ons. Cancers (Basel), 2025. 17(13).

4. Rizwani, F., P. PaJl, and K. Jain, Unlocking glioblastoma: breakthroughs in molecular mechanisms and next-genera*on therapies. Med Oncol, 2025. 42(7): p. 276.

5. Guo, X., et al., Neuronal Ac*vity Promotes Glioma Progression by Inducing Proneural-to-Mesenchymal Transi*on in Glioma Stem Cells. Cancer Res, 2024. 84(3): p. 372–387.

6. Johung, T. and M. Monje, Neuronal ac*vity in the glioma microenvironment. Curr Opin Neurobiol, 2017. 47: p. 156–161.

7. !!! INVALID CITATION !!!.

8. Gillespie, S. and M. Monje, An ac*ve role for neurons in glioma progression: making sense of Scherer’s structures. Neuro Oncol, 2018. 20(10): p. 1292–1299.

9. De Silva, M.I., B.W. Stringer, and C. Bardy, Neuronal and tumourigenic boundaries of glioblastoma plas*city. Trends Cancer, 2023. 9(3): p. 223–236.

10. Krishna, S., et al., Glioblastoma remodelling of human neural circuits decreases survival. Nature, 2023. 617(7961): p. 599-607.

11. Montgomery, M.K., et al., Glioma-Induced Altera*ons in Neuronal Ac*vity and Neurovascular Coupling during Disease Progression. Cell Rep, 2020. 31(2): p. 107500.

12. Pei, Z., et al., Pathway analysis of glutamate-mediated, calcium-related signaling in glioma progression. Biochem Pharmacol, 2020. 176: p. 113814.

13. Sontheimer, H., A role for glutamate in growth and invasion of primary brain tumors. J Neurochem, 2008. 105(2): p. 287–95.

14. Venkatesh, H.S., et al., Electrical and synap*c integra*on of glioma into neural circuits. Nature, 2019. 573(7775): p. 539-545.

15. Piao, Y., et al., Glioblastoma resistance to an*-VEGF therapy is associated with myeloid cell infiltra*on, stem cell accumula*on, and a mesenchymal phenotype. Neuro Oncol, 2012. 14(11): p. 1379–92.

16. Wild-Bode, C., et al., Sublethal irradia*on promotes migra*on and invasiveness of glioma cells: implica*ons for radiotherapy of human glioblastoma. Cancer Res, 2001. 61(6): p. 2744–50.

17. Sprugnoli, G., A.J. Golby, and E. Santarnecchi, Newly discovered neuron-to-glioma communica*on: new noninvasive therapeu*c opportuni*es on the horizon? Neurooncol Adv, 2021. 3(1): p. vdab018.

18. Venkatesh, H.S., et al., Targe*ng neuronal ac*vity-regulated neuroligin-3 dependency in high-grade glioma. Nature, 2017. 549(7673): p. 533–537.

19. Venkatesh, H.S., Targe*ng electrochemical communica*on between neurons and cancer. Sci Transl Med, 2023. 15(706): p. eadi5170.

20. Smirnova, L. and T. Hartung, The Promise and Poten*al of Brain Organoids. Adv Healthc Mater, 2024. 13(21): p. e2302745.

21. Liu, Y., et al., Pa*ent-derived xenograft models in cancer therapy: technologies and applica*ons. Signal Transduct Target Ther, 2023. 8(1): p. 160.

22. Yadav, N. and B.W. Purow, Understanding current experimental models of glioblastoma-brain microenvironment interac*ons. J Neurooncol, 2024. 166(2): p. 213–229.

23. Varn, F.S., et al., Glioma progression is shaped by gene*c evolu*on and microenvironment interac*ons. Cell, 2022. 185(12): p. 2184–2199. e16.

24. Mancusi, R. and M. Monje, The neuroscience of cancer. Nature, 2023. 618(7965): p. 467–479.

25. Silverman, D.A., et al., Cancer-associated neurogenesis and nerve-cancer cross-talk. Cancer research, 2021. 81(6): p. 1431–1440.

26. Fernandopulle, M.S., et al., Transcrip*on factor–mediated differen*a*on of human iPSCs into neurons. Current protocols in cell biology, 2018. 79(1): p. e51.

27. Le, N.T., et al., Prion protein pathology in Ubiquilin 2 models of ALS. Neurobiol Dis, 2024. 201: p. 106674.

28. Kaech, S. and G. Banker, Culturing hippocampal neurons. Nat Protoc, 2006. 1(5): p. 2406–15.

29. Seker-Polat, F., et al., Tumor Cell Infiltra*on into the Brain in Glioblastoma: From Mechanisms to Clinical Perspec*ves. Cancers (Basel), 2022. 14(2).

30. Allen, M., et al., Origin of the U87MG glioma cell line: Good news and bad news. Sci Transl Med, 2016. 8(354): p. 354re3.

31. Tallman, M.M., et al., The small molecule drug CBL0137 increases the level of DNA damage and the efficacy of radiotherapy for glioblastoma. Cancer Led, 2021. 499: p. 232–242.

32. Gorai, S., G. Rathore, and K. Das, Selume*nib-A Comprehensive Review of the New FDA-Approved Drug for Neurofibromatosis. Indian Dermatol Online J, 2024. 15(4): p. 701–705.

33. Kelemen, B.R., K. Hsiao, and S.A. Goueli, Selec*ve in vivo inhibi*on of mitogen-ac*vated protein kinase ac*va*on using cell-permeable pep*des. J Biol Chem, 2002. 277(10): p. 8741–8.

34. Venkataramani, V., et al., Glioblastoma hijacks neuronal mechanisms for brain invasion. Cell, 2022. 185(16): p. 2899–2917.e31.

35. Tetzlaff, S.K., et al., Characterizing and targe*ng glioblastoma neuron-tumor networks with retrograde tracing. Cell, 2025. 188(2): p. 390–411.e36.

36. Zeng, Y., et al., The Tumorgenicity of Glioblastoma Cell Line U87MG Decreased During Serial In Vitro Passage. Cell Mol Neurobiol, 2018. 38(6): p. 1245–1252.

37. Zeng, X., et al., The expression of G protein-coupled receptor kinase 5 and its interac*on with dendri*c marker microtubule-associated protein-2 a[er status epilep*cus. Epilepsy Research, 2017. 138: p. 62–70.

38. Venkataramani, V., et al., Glutamatergic synap*c input to glioma cells drives brain tumour progression. Nature, 2019. 573(7775): p. 532–538.

39. Zeng, Q., et al., Synap*c proximity enables NMDAR signalling to promote brain metastasis. Nature, 2019. 573(7775): p. 526–531.

40. Corrêa, S.A. and K.L. Eales, The Role of p38 MAPK and Its Substrates in Neuronal Plas*city and Neurodegenera*ve Disease. J Signal Transduct, 2012. 2012: p. 649079.

41. Tallman, M., et al., The small molecule drug CBL0137 increases the level of DNA damage and the efficacy of radiotherapy for glioblastoma. Cancer Leders, 2020. 499.

42. Tallman, M.M., et al., Improving Localized Radiotherapy for Glioblastoma via Small Molecule Inhibi*on of KIF11. Cancers (Basel), 2023. 15(12).

43. Wang, C., et al., Scalable produc*on of iPSC-derived human neurons to iden*fy tau-lowering compounds by high-content screening. Stem cell reports, 2017. 9(4): p. 1221–1233.

44. Fang, C., et al., Prions ac*vate a p38 MAPK synaptotoxic signaling pathway. PLoS Pathog, 2018. 14(9): p. e1007283.

45. Le, N.T.T., B. Wu, and D.A. Harris, Prion neurotoxicity. Brain Pathol, 2019. 29(2): p. 263–277.

46. Srivastava, D.P., K.M. Woolfrey, and P. Penzes, Analysis of dendri*c spine morphology in cultured CNS neurons. J Vis Exp, 2011(53): p. e2794.

47. Zhou, Y., et al., Metascape provides a biologist-oriented resource for the analysis of systems-level datasets. Nat Commun, 2019. 10(1): p. 1523.

48. Shannon, P., et al., Cytoscape: a so[ware environment for integrated models of biomolecular interac*on networks. Genome Res, 2003. 13(11): p. 2498–504.

