## Supplemental Data for "A Human Neuronal Co-Culture System Reveals Early Tumor-Neuron Communication and Targetable Pathways in Glioblastoma"

**Figure S1. Synaptic abnormalities in neurons in response to glioblastoma-derived signals.**

Neurons from dual-interface co-cultures were analyzed for synaptic protein expression across three groups: control neurons (Ctrl-neuron), neurons exposed to U-87MG (U-87MG-neuron), and neurons exposed to NU-757 (NU-757-neuron).

**(A-B)** Western blot analysis was performed to assess the expression of synaptophysin (presynaptic marker), PSD-95 and NMDAR2B (postsynaptic markers), with actin used as a loading control. Representative immunoblots (A) and corresponding densitometric quantification (B) revealed a significant reduction in PSD-95 and NMDAR2B expression in neurons exposed to U-87MG and NU-757, indicating impaired postsynaptic signaling. Data are presented as mean ± SEM. Significance: ns (not significant), **P < 0.01, ***P < 0.001, ****P < 0.0001, exposed neurons vs. control neurons, using unpaired t-tests.

**(C–D)** Neuronal MAP-2 (Microtubule-Associated Protein 2) expression was assessed in control and U-87 MG-exposed neurons. Representative immunofluorescence images (C) showed markedly reduced MAP-2 signal (green) in glioblastoma-exposed neurons, consistent with western blot results (D). Data represent 18-21 neuron segments. Values are shown as mean ± SD. Unpaired t-tests were used for statistical analysis; *P < 0.05. (D) Western blots were probed for MAP-2, with actin as a loading control. Quantitative densitometric analysis confirmed a significant decrease in MAP-2 expression in glioblastoma-exposed neurons. Data represent three independent experiments (n = 3) and are presented as mean ± SEM. **P < 0.01, exposed neurons vs. control, using unpaired t-tests.

**Figure S2. Quantitative proteomics highlights differentially regulated proteins in neurons exposed to glioblastoma.**

Quantitative proteomic analysis was performed on neurons exposed to conditioned media from U-87MG (Neurons_U-87), NU-757 (Neurons_NU-757), and untreated control neurons (n = 4 per group).

(A) Venn diagram showing the total number of proteins detected in Neurons_U-87 and Neurons_NU-757. More than 5,000 proteins were identified in each group, with 4,965 proteins shared, indicating a highly overlapping baseline proteome.

(B) Differential expression analysis using a fold change threshold of >1.5 or <0.5 revealed 91 and 520 differentially expressed proteins in Neurons_U-87 and Neurons_NU-757, respectively. Only 8 proteins were commonly dysregulated, suggesting distinct proteomic responses to each glioblastoma line.

(C) Venn diagram of significantly dysregulated proteins ($p < 0.05$) showing 36 proteins commonly altered in both Neurons_U-87 and Neurons_NU-757, indicating a partially shared but predominantly cell type-specific neuronal response to glioblastoma-derived factors.

**Figure S3. Distinct proteomic profiles and functional enrichments in neurons exposed to patient-derived versus standard glioblastoma cells.**

**(A)** Network visualization of enriched functional terms from differentially expressed proteins in neurons exposed to NU-757 versus U-87 MG glioblastoma cells for 24 hours.

Nodes are colored by cluster ID, with nodes sharing the same ID positioned near each other to indicate functional similarity.

**(B)** The same enrichment network as in (A), represented using pie charts. Each node is color-coded by the contributing protein list: NU-757 (red) or U-87 MG (blue). **(C-D)** Volcano plots of differentially expressed proteins in neurons exposed to U-87 MG (C) and NU-757 (D) glioblastoma cells compared to control neurons. Proteins are plotted by statistical significance ($-\log_{10}P$ value, y-axis) and fold change ($\log_2$ FC, x-axis). Significance thresholds are set at fold change (FC) ≥ 1.3 and $P < 0.05$. Red dots indicate significantly upregulated proteins, blue dots indicate significantly downregulated proteins, and grey dots indicate proteins with no significant change.

**Figure S4. Alteration of MLK2 and p38 MAPK signaling pathways in neurons exposed to glioblastoma cells.**

**(A-B)** Immunoblot analysis of neuronal lysates from dual-interface co-cultures, including control neurons (Ctrl-neuron), neurons exposed to U-87 MG glioblastoma cells (U-87 MG-neuron), and neurons exposed to patient-derived NU-757 glioblastoma cells (NU-757-neuron). Blots were probed for MLK2, total p38, phosphorylated p38 (p-p38, Thr180/Tyr182), and actin as a loading control. Data represent three independent experiments (n = 3). Lower panel (B) showed the densitometric quantification of western blot images. Values are presented as mean ± SEM. Statistical comparisons were performed using unpaired t-tests. Significance is indicated as: ns (not significant), *P < 0.05, ****P < 0.0001.

**(C)** Representative triple immunofluorescence images showing phalloidin (gray), p38 (green), and MLK2 (red) in neurons following 24-hour exposure to glioblastoma. DAPI (blue) marks nuclei. Dotted lines delineate nuclear and dendritic spine regions based on phalloidin and DAPI staining. Scale bars = 10 μm (nuclear panels) and 2 μm (spine panels).

**(D-E)** Quantification of p38 (D) and MLK2 (E) immunofluorescence intensity in neuronal nuclei and dendritic spines from (C). Data were pooled from 62-82 nuclear regions and 997-2163 spine regions across three independent experiments. Values are presented as mean ± SEM. Statistical comparisons were performed using unpaired t-tests. Significance is indicated as: ns (not significant), ****P < 0.0001.

**Figure S5. PP2A involvement in ERK signaling and differential drug responses in glioblastoma models.**

**(A)** Protein–protein interaction (PPI) network centered on ERK1/2 (MAPK3) and PP2A subunits (PPP2R1B and PPP2R5B), showing significantly altered proteins in neurons exposed to U-87 MG glioblastoma.

**(B)** Immunoblot analysis of neuronal lysates from dual-interface co-cultures, including control neurons (Ctrl-neuron) and neurons exposed to U-87 MG glioblastoma (U-87 MG-neuron). Blots were probed for PP2A B subunit and PPP2R5B and actin (loading control). Data represent three independent experiments (n = 3). Lower panel showed the densitometric quantification of western blot images. Values are presented as mean ± SEM. Statistical comparisons were performed using unpaired t-tests. Significance is indicated as: ns (not significant), *P < 0.05.

**(C-D)** Quantification of glioblastoma cell proliferation using Incucyte phase-contrast imaging. Selumetinib (MEK inhibitor) reduced proliferation in U-87 MG cells (C) but had no significant effect on NU-757 cells (D). Values are shown as mean ± SEM. Data were pooled from 4 independent cultures per condition, with 5 fields imaged per culture. **(E)** Barrier membrane blocks cell migration. Validation of the membrane's inhibitory efficacy was performed by comparing the non-invasive HMC3 microglial cell line with the invasive U-87 MG glioblastoma cell line, confirming selective prevention of tumor cell migration. (**F**) Migration assay of U-87 MG glioblastoma cells using Incucyte phase-contrast imaging. Selumetinib treatment inhibited both proliferation and migration of U-87 MG cells. Values are shown as mean ± SEM. Data were pooled from 3 independent cultures per condition, with 25 fields imaged per culture.

**Figure S6. Effects of PP2A activation and combined MEK inhibition on glioblastoma and microglial cell proliferation.**

**(A-B)** Quantification of cell proliferation using Incucyte phase-contrast imaging in U-87 MG glioblastoma cells (A) and HMC3 microglia (B) treated with DT-061 (PP2A activator) at increasing concentrations. DT-061 had minimal effects on both cell types, except at 10 μM, where it moderately reduced U-87 MG proliferation.

**(C-D)** Proliferation assays of U-87 MG glioblastoma (C) and HMC3 microglia (D) treated with a combination of Selumetinib (MEK inhibitor) and DT-061. The combination significantly suppressed U-87 MG cell proliferation, with partial inhibitory effects observed in HMC3 cells at 10 μM DT-061.

**(E)** Neurons treated with the specified inhibitors showed no signs of toxicity and maintained healthy morphology over a 4-day culture period.

Values are presented as mean ± SEM. Data were pooled from 4 independent cultures per condition, with 5 fields imaged per culture. Ctrl = untreated control; Sel = Selumetinib; DT = DT-061.

**Figure S1.**

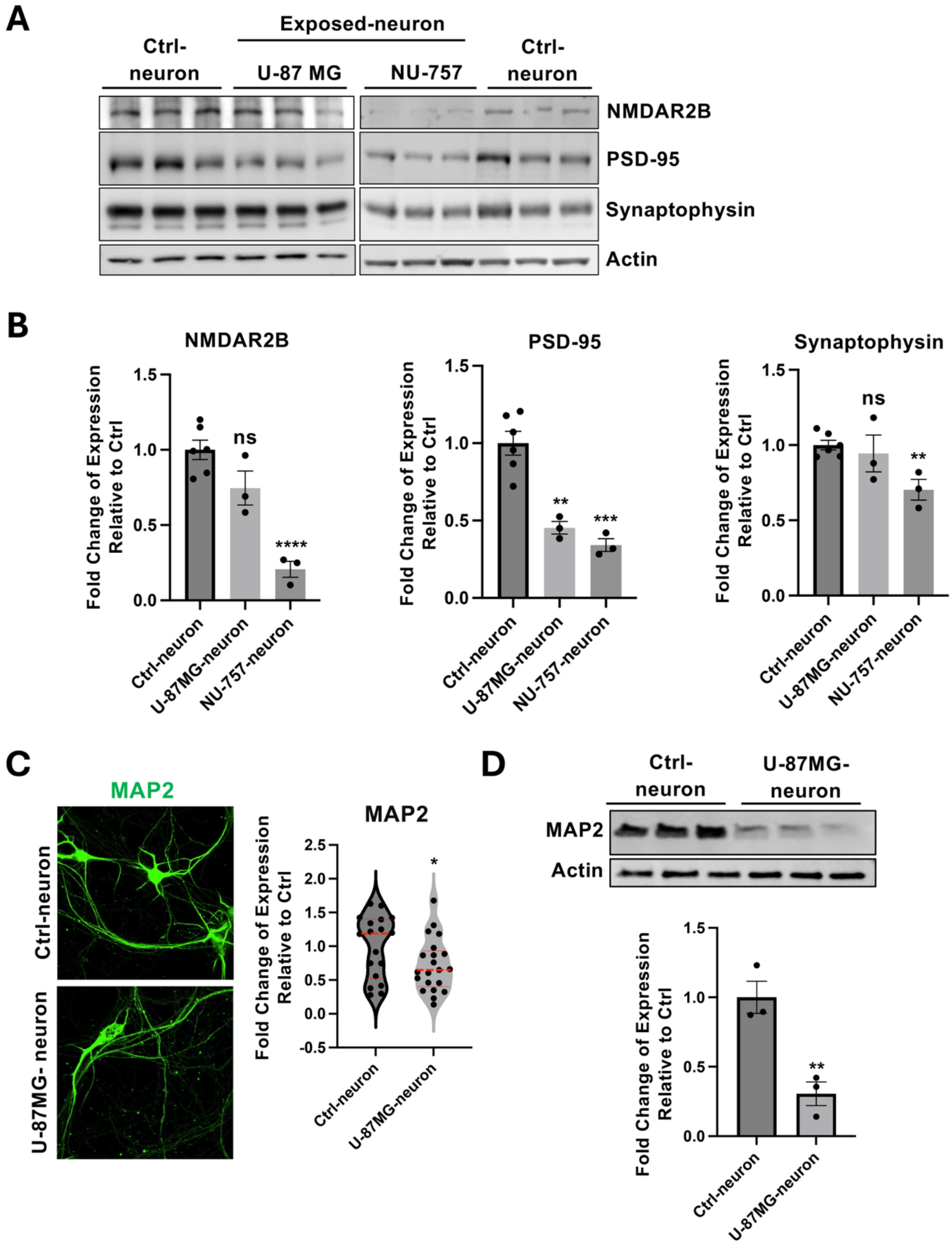
A
Exposed-neuron
Ctrl-neuron
U-87 MG
NU-757
Ctrl-neuron
NMDAR2B
PSD-95
Synaptophysin
Actin
B
NMDAR2B
PSD-95
Synaptophysin
Fold Change of Expression Relative to Ctrl
ns
****
**
***
**
Ctrl-neuron
U-87MG-neuron
NU-757-neuron
C
MAP2
Ctrl-neuron
U-87MG- neuron
*
D
Ctrl-neuron
U-87MG-neuron
MAP2
Actin

**Figure S2.**

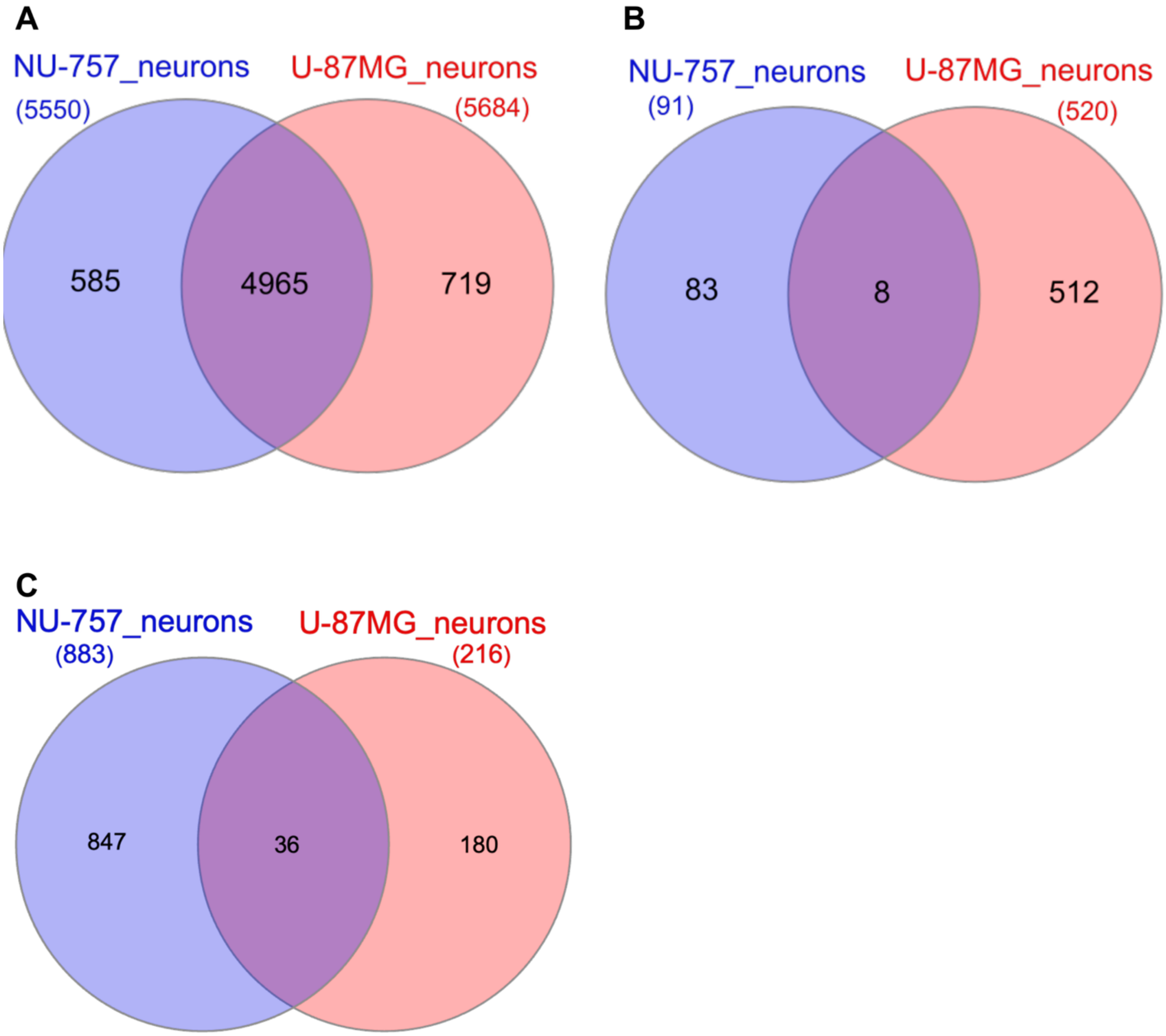
A
NU-757_neurons
(5550)
U-87MG_neurons
(5684)
585
4965
719
B
NU-757_neurons
(91)
U-87MG_neurons
(520)
83
8
512
C
NU-757_neurons
(883)
U-87MG_neurons
(216)
847
36
180

### Figure S3.

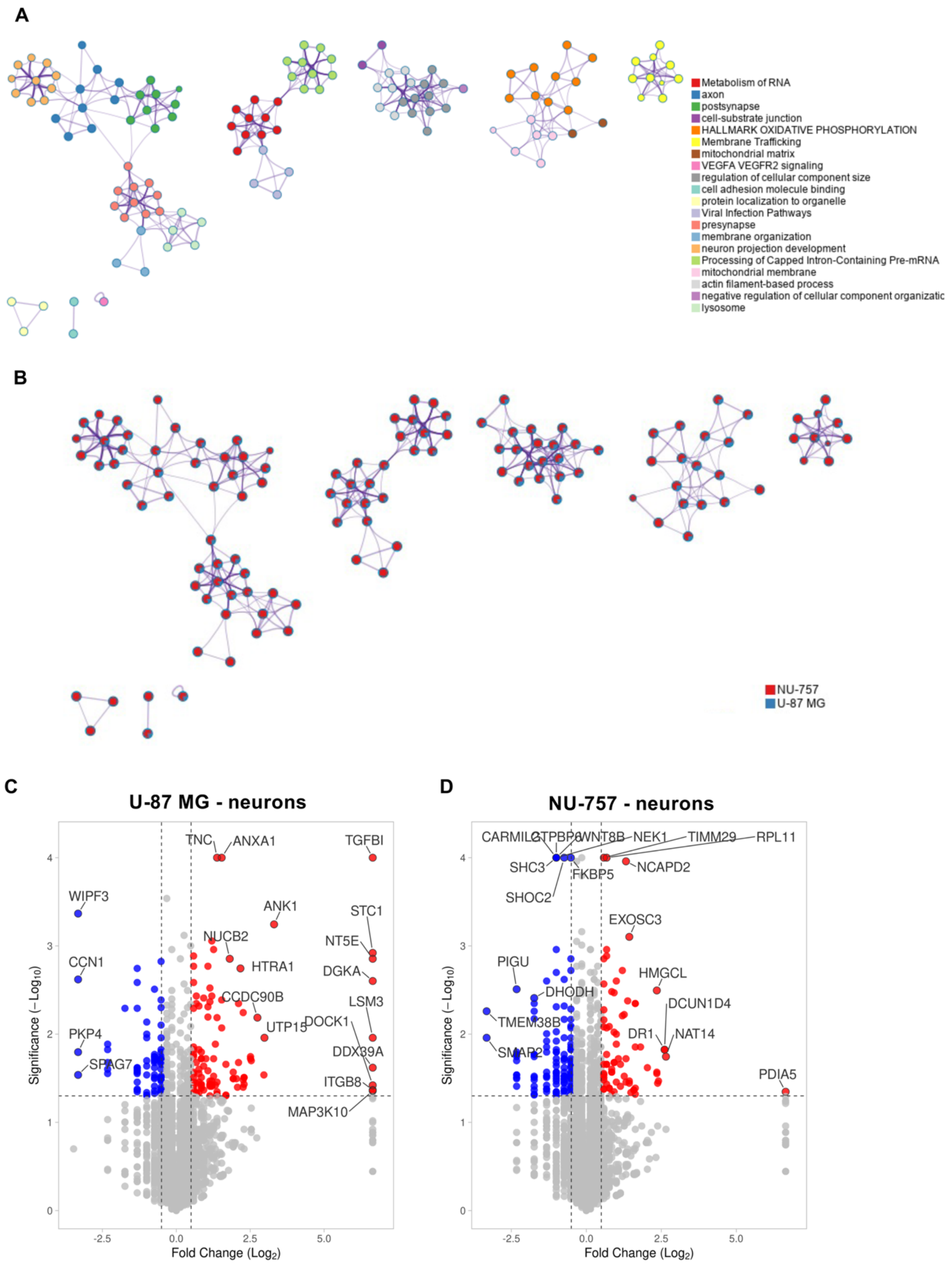
A
Metabolism of RNA
axon
postsynapse
cell-substrate junction
HALLMARK OXIDATIVE PHOSPHORYLATION
Membrane Trafficking
mitochondrial matrix
VEGFA VEGFR2 signaling
regulation of cellular component size
cell adhesion molecule binding
protein localization to organelle
Viral Infection Pathways
presynapse
membrane organization
neuron projection development
Processing of Capped Intron-Containing Pre-mRNA
mitochondrial membrane
actin filament-based process
negative regulation of cellular component organizatic
lysosome
B
NU-757
U-87 MG
C
U-87 MG - neurons
TNC
ANXA1
TGFBI
WIPF3
ANK1
STC1
NUCB2
NT5E
CCN1
HTRA1
DGKA
CCDC90B
LSM3
PKP4
UTP15
DOCK1
SPAG7
DDX39A
ITGB8
MAP3K10
Significance (−Log10)
Fold Change (Log2)
D
NU-757 - neurons
CARMIL2
GTPBP6
WNT8B
NEK1
TIMM29
RPL11
SHC3
NCAPD2
FKBP5
SHOC2
EXOSC3
PIGU
HMGCL
DHODH
DCUN1D4
TMEM38B
DR1
NAT14
SMAP2
PDIA5
Significance (−Log10)
Fold Change (Log2)

**Figure S4.**

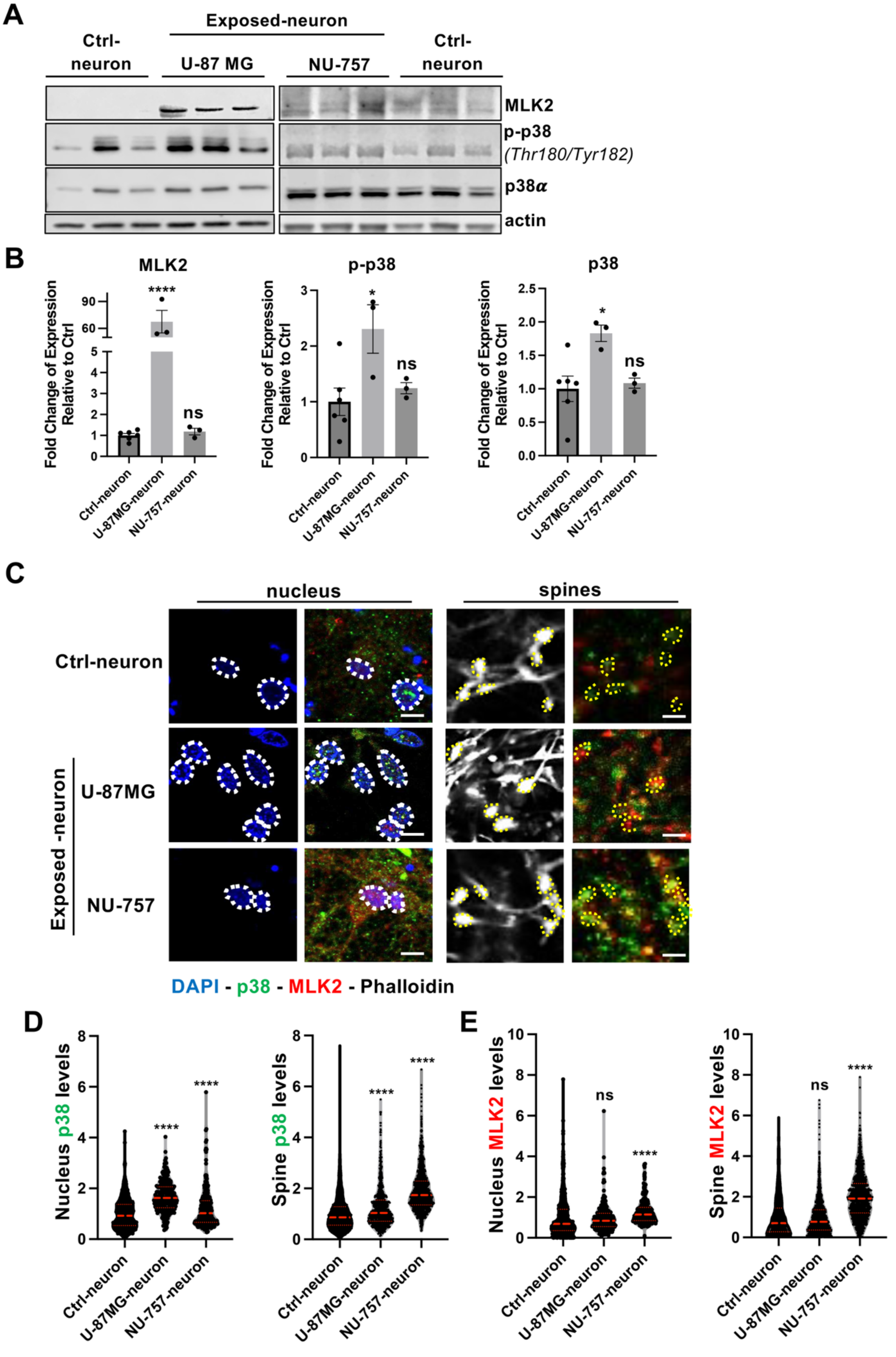
A
Exposed-neuron
Ctrl-neuron
U-87 MG
NU-757
Ctrl-neuron
MLK2
p-p38
(Thr180/Tyr182)
p38α
actin
B
MLK2
p-p38
p38
Fold Change of Expression Relative to Ctrl
Ctrl-neuron
U-87MG-neuron
NU-757-neuron
****
*
*
ns
C
nucleus
spines
Ctrl-neuron
Exposed -neuron
U-87MG
NU-757
DAPI - p38 - MLK2 - Phalloidin
D
Nucleus p38 levels
Spine p38 levels
E
Nucleus MLK2 levels
Spine MLK2 levels

**Figure S5.**

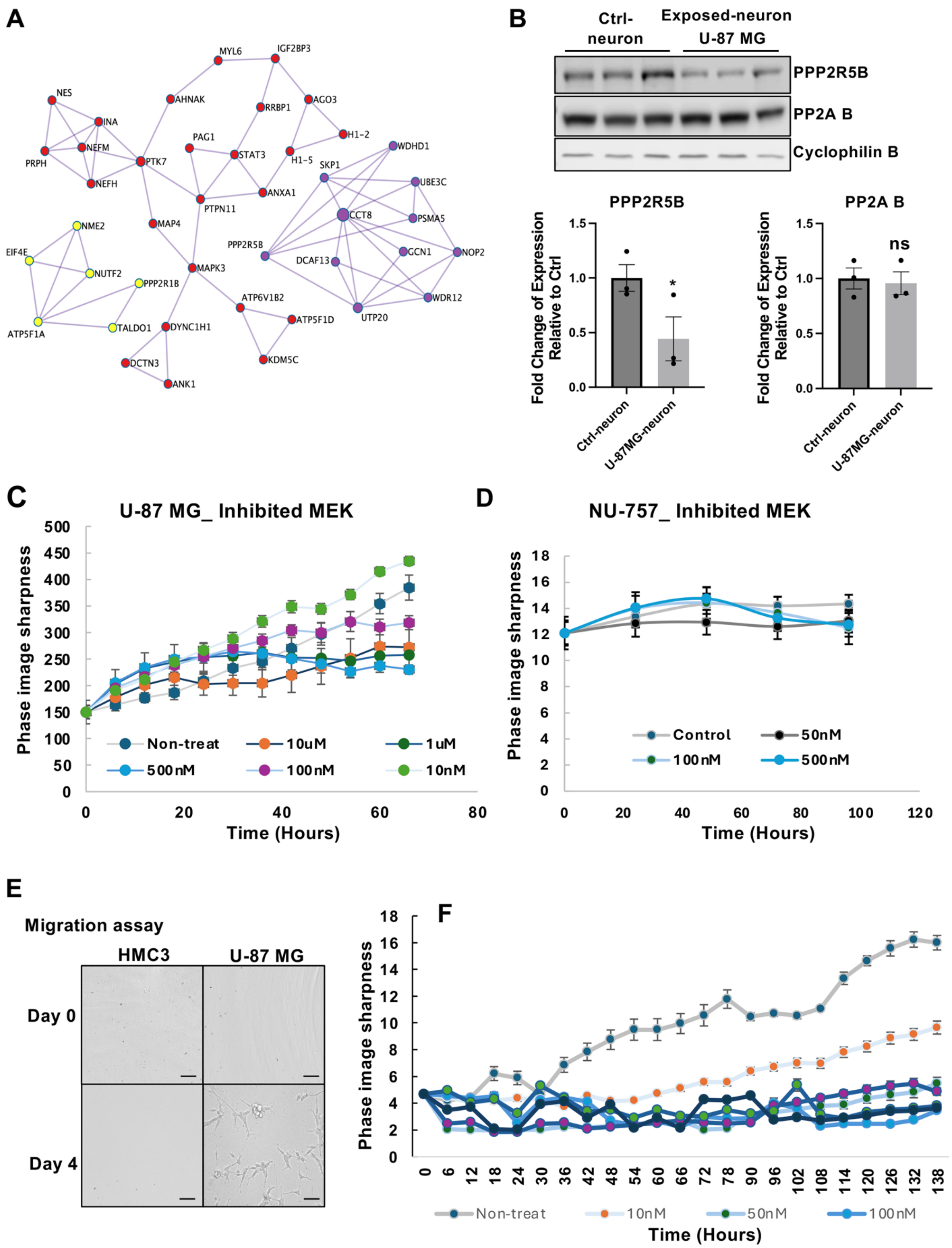
A
NES
INA
NEFM
PRPH
NEFH
PTK7
AHNAK
MYL6
IGF2BP3
RRBP1
AGO3
H1-2
H1-5
PAG1
STAT3
PTPN11
ANXA1
MAP4
MAPK3
DYNC1H1
DCTN3
ANK1
ATP6V1B2
ATP5F1D
KDM5C
NME2
EIF4E
NUTF2
PPP2R1B
TALDO1
ATP5F1A
WDHD1
SKP1
UBE3C
CCT8
PSMA5
PPP2R5B
DCAF13
GCN1
NOP2
WDR12
UTP20
B
Ctrl-neuron
Exposed-neuron U-87 MG
PPP2R5B
PP2A B
Cyclophilin B
PPP2R5B
PP2A B
Fold Change of Expression Relative to Ctrl
Ctrl-neuron
U-87MG-neuron
*
ns
C
U-87 MG_ Inhibited MEK
Phase image sharpness
Time (Hours)
Non-treat
10uM
1uM
500nM
100nM
10nM
D
NU-757_ Inhibited MEK
Phase image sharpness
Time (Hours)
Control
50nM
100nM
500nM
E
Migration assay
HMC3
U-87 MG
Day 0
Day 4
F
Phase image sharpness
Time (Hours)
Non-treat
10nM
50nM
100nM

Figure S6.

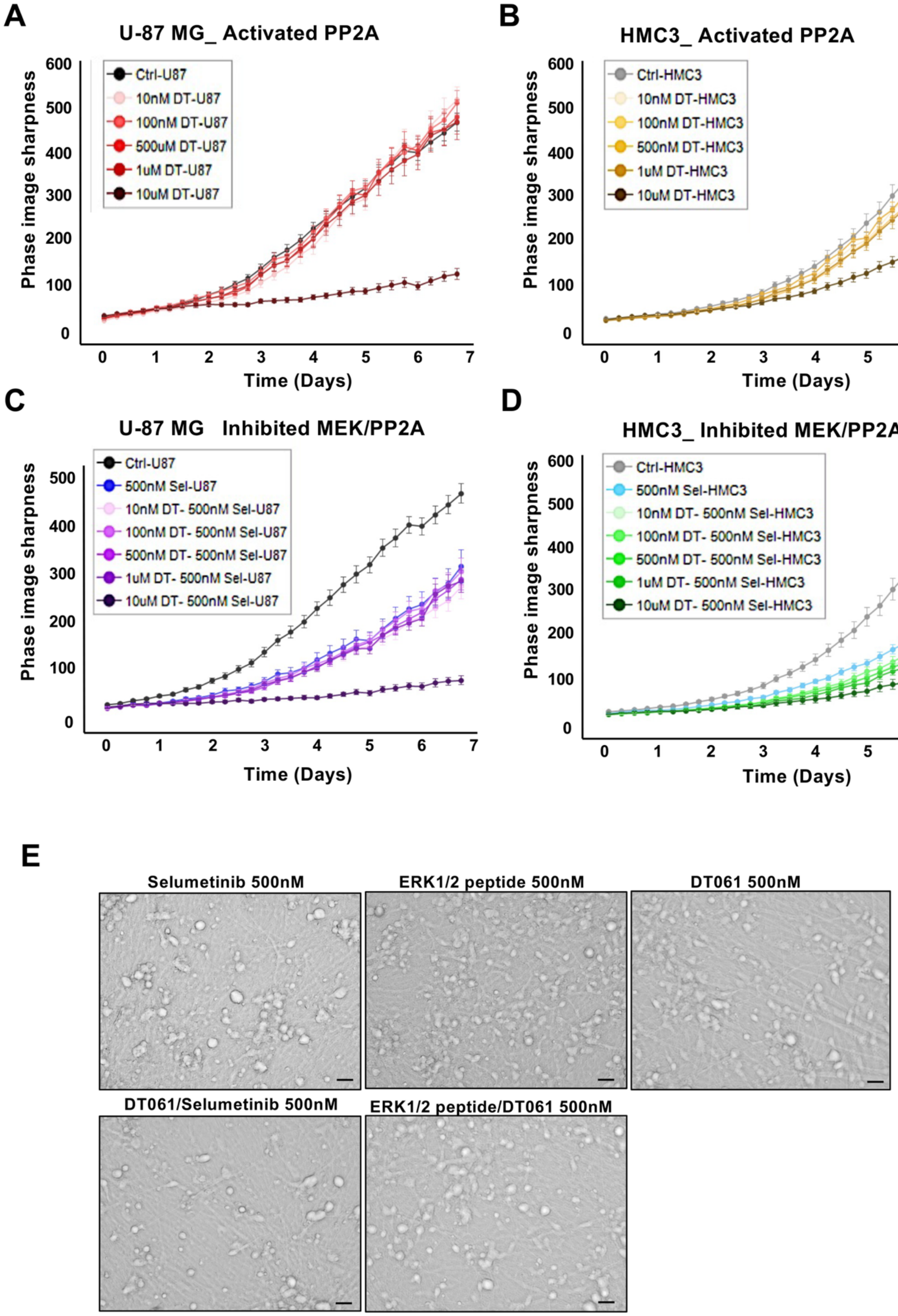

A
U-87 MG_ Activated PP2A
Ctrl-U87
10nM DT-U87
100nM DT-U87
500uM DT-U87
1uM DT-U87
10uM DT-U87
Phase image sharpness
Time (Days)
B
HMC3_ Activated PP2A
Ctrl-HMC3
10nM DT-HMC3
100nM DT-HMC3
500nM DT-HMC3
1uM DT-HMC3
10uM DT-HMC3
Phase image sharpness
Time (Days)
C
U-87 MG Inhibited MEK/PP2A
Ctrl-U87
500nM Sel-U87
10nM DT- 500nM Sel-U87
100nM DT- 500nM Sel-U87
500nM DT- 500nM Sel-U87
1uM DT- 500nM Sel-U87
10uM DT- 500nM Sel-U87
Phase image sharpness
Time (Days)
D
HMC3_ Inhibited MEK/PP2A
Ctrl-HMC3
500nM Sel-HMC3
10nM DT- 500nM Sel-HMC3
100nM DT- 500nM Sel-HMC3
500nM DT- 500nM Sel-HMC3
1uM DT- 500nM Sel-HMC3
10uM DT- 500nM Sel-HMC3
Phase image sharpness
Time (Days)
E
Selumetinib 500nM
ERK1/2 peptide 500nM
DT061 500nM
DT061/Selumetinib 500nM
ERK1/2 peptide/DT061 500nM

**Table S1:**

List of antibodies and dyes used in the study

| Antibody/Dye target | Species | Dilution | | Company (Catalog #) |
|---|---|---|---|---|
| | | IF | WB | |
| Anti-ERK1/2 (ERK-7D8) | Mouse | 1:200 | 1:1000 | Invitrogen (13-6200) |
| Anti-Phospho-ERK1 (T202/Y204)/ERK2 (T185/Y187) | Rabbit | 1:200 | 1:1000 | R&D Systems (MAB1018) |
| Anti -p38α MAPK | Rabbit | 1:200 | 1/1000 | Cell Signaling (#9218) |
| Anti-Phospho-p38 MAPK (Thr180/Tyr182) | Rabbit | 1:100 | 1:1000 | Cell Signaling (#9211) |
| Anti-Map2 | Rabbit | 1:200 | 1:1000 | Abcam (ab32454) |
| Anti-MLK2 | Rabbit | 1:200 | 1:1000 | Invitrogen PA5-75848 |
| Anti-PPP2R5B | Rabbit | 1:100 | 1:1000 | Abcam (ab181023) |
| Rhodamine-Phalloidin | - | 1:500 | - | ThermoFisher (R415) |
| Alexa Fluor 488-phalloidin | - | 1:500 | - | ThermoFisher (A12379) |
| Anti-NMDAR2B | Mouse | - | 1:1000 | Abcam (ab93610) |
| Synaptophysin | Mouse | - | 1:1000 | Santa Cruz Biotechnology (sc-17750) |
| Anti-PSD95 | Mouse | - | 1:1000 | Abcam (ab18258)) |
| PP2A B Subunit (2G9) | Mouse | - | 1:1000 | Cell Signaling (mAb #5689) |
| Anti-PPP2R5B | Rabbit | - | 1:1000 | Antibodies-online (ABIN6243837) |
| Anti-β-Actin | Mouse | - | 1:2000 | Cell Signaling (#4967) |
| anti-Cyclophilin B (D1V5J) | Rabbit | - | 1:2000 | Cell Signaling (#43603) |
| Anti-rabbit IgG, HRP-linked Antibody | Rabbit | - | 1:2000 | Cell Signaling (#7074**)** |
| Anti-mouse IgG, HRP-linked Antibody | Mouse | - | 1:2000 | Cell Signaling (#7076) |
| Anti-β-Actin | Mouse | - | 1:2000 | Cell Signaling (#4967) |
| DAPI | - | 1/5000 | | Invitrogen(D1306) |
| Alexa Flour 488 | Mouse | 1:200 | - | Invitrogen (A32766) |
| Alexa Flour 647 | Rabbit | 1:200 | - | Invitrogen (A-32733) |
| IRDye 680DR | Rabbit | | 1:10000 | LI-COR (926-68073) |
| IRDye 800CW | Mouse | | 1:10000 | LI-COR (926-32212) |
